# A double-hit of social and economic stress in mice precipitates changes in decision-making strategies

**DOI:** 10.1101/2023.03.19.533304

**Authors:** Romain Durand-de Cuttoli, Freddyson J. Martínez-Rivera, Long Li, Angélica Minier-Toribio, Zhe Dong, Denise J. Cai, Scott J. Russo, Eric J. Nestler, Brian M. Sweis

## Abstract

Economic stress can serve as a “second-hit” for those who already accumulated a history of adverse life experiences. How one recovers from a setback is a core feature of resilience but is seldom captured in animal studies. We challenged mice in a novel two-hit stress model by exposing animals to chronic social defeat stress (first-hit) and then testing how mice adapt to reward scarcity on a neuroeconomic task (second-hit). Mice were tested longitudinally across months on the Restaurant Row task during which mice foraged daily for their sole source of food while on a limited time budget. An abrupt transition into a reward-scarce environment on this task elicits an economic crisis, precipitating a massive drop in food intake and body weight to which mice must respond in order to survive. We found that mice with a history of social defeat mounted a robust behavioral response to this economic challenge. This recovery was achieved through a complex redistribution of how time was allocated among competing opportunities via multiple valuation algorithms. Interestingly, we found that mice with a history of social defeat displayed changes in the development of decision-making policies during the recovery process important for not only ensuring food security necessary for survival but also prioritizing subjective value. These findings indicate that an individual’s capacity to “bounce back” from economic stress depends on one’s prior history of stress and can affect multiple aspects of subjective well-being, highlighting a motivational balance that may be altered in stress-related disorders such as depression.

**In Brief:** Durand-de Cuttoli et al. found that after chronic social defeat stress, when mice were subsequently challenged on a neuroeconomic foraging task, an economic stressor can serve as a “second hit” and reveal changes in the development of complex decision-making strategies important for maintaining the balance between food security and subjective well-being.

---

Repeated exposure to stress can drive maladaptive behaviors and contribute to mental illness.^1^ How one responds to multiple bouts of stress, including how one recovers back to baseline following stress, is critical for generating healthy coping strategies and can separate those who are resilient from those who might be susceptible to developing psychiatric sequelae.^2-4^ In the animal stress literature, studies of the behavioral and neurobiological underpinnings of resilience have gained traction in recent years.^5,6^ However, studies of resilience have only sparsely focused on characterizing the behavioral and cognitive processes involved in one’s journey during recovery to baseline levels, partly because operationalizing a model capable of capturing a stress-inducing setback and associated signatures of recovery has been difficult to develop in animals.

A unique way to formalize and capture a stress-inducing setback in a translationally relevant manner involves turning to the neuroeconomics decision-making field. Neuroeconomics describes the study of how the physical limits of the brain give rise to cognitive mechanisms involved in decision-making information processing.^7,8^ This field focuses on understanding complex interactions between multiple choice parameters including reward value, price, effort expenditure, energetic demand, competing action-selection processes, and opportunity costs.^9^ These factors can differentially influence fundamentally distinct valuation algorithms served by dissociable decision processes in physically separable circuits in the brain.^10^ In this context, an example of economic stress can be defined as a change in one’s budget constraints, (e.g., in humans, the financial burden of losing one’s job or following market inflation), which can take a cognitive and affective toll on an individual in addition to the practical ramifications a stricter budget can have.^11,12^ A stressful setback such as this can precipitate changes in behavior that may attempt to correct or react to one’s environment and may be adaptive or maladaptive in nature. By studying stress in this way, more robust measures of decision-making related behavior can be gleaned from how one responds to economic challenges. Modeling this type of stressor has received far less attention in the animal stress literature. For animal research, approaches in neuroeconomics offer a novel framework with which to operationalize these concepts in ways that may be useful for understanding stress-related pathologies with richer behavioral endpoints.^13^

We set out to examine how an individual’s prior history of stress influences future responses to stress. We investigated exposure to two distinct but commonly interacting types of stress often highlighted in human but not animal work: social stress and economic stress.^14-16^ Here, we developed a novel “two-hit” stress model by combining the well-established chronic social defeat stress protocol^17^ (first hit) with a longitudinal neuroeconomic decision-making paradigm, “Restaurant Row.”^18-20^ This complex task, during which mice must forage for their sole source of food in a limited period of time, is comprised of a changing economic landscape that embeds an acute and striking economic challenge (second hit). This economic setback precipitates a crisis to which mice must respond for survival. This approach enabled us to rapidly extract individual differences in response to social stress and investigate how these different profiles affect the ways in which subsequent economic stress alters the balance of multiple decision-making strategies during the recovery process.

## Methods

### Subjects

Adult male C57BL/6J mice (Jackson, 10wks, n=32) and CD-1 mice (Charles River, 20wks, n=22) were used. After social defeat, mice were individually housed and maintained on a 12-hr light/dark cycle with ad libitum water, only food-restricted during Restaurant Row testing, conducted during the light phase. Experiments were approved by the Mount Sinai Institutional Animal Care and Use Committee (IACUC; protocol number LA12-00051) and adhered to the National Institutes of Health (NIH) guidelines.

### Chronic Social Defeat Stress

22 C57BL/6J mice underwent chronic social defeat (Fig. 1a, Fig. S1a). Each mouse was paired with a CD-1 aggressor mouse and physically interacted for 5 min before remaining co-housed separated by a mesh divider for the rest of the day. This was repeated with a novel CD-1 mouse across 10 days. 10 additional C57BL/6J non-defeated mice were paired instead with other C57BL/6J mice. Social avoidance induced by this protocol (Fig. S1b, *F*_1,31_=10.793, *p*<0.01) has served as a well-validated predictor of several additional depression-related phenotypes on other rapid behavioral screening tests.^17^ The social interaction test is a short procedure during which a single C57BL/6J mouse is placed in a large open field arena with a novel CD-1 mouse enclosed in a small chamber. EthoVision software was used to track the location of the C57BL/6J mouse. Time spent near (interaction zone) versus away from the CD-1 mouse relative to when no CD-1 mouse is present is used to calculate a social interaction score to split mice into resilient and susceptible subgroups. Mice were then food restricted to 80-85% of their free-feeding body weight over the next 3 days.

**Figure 1.**
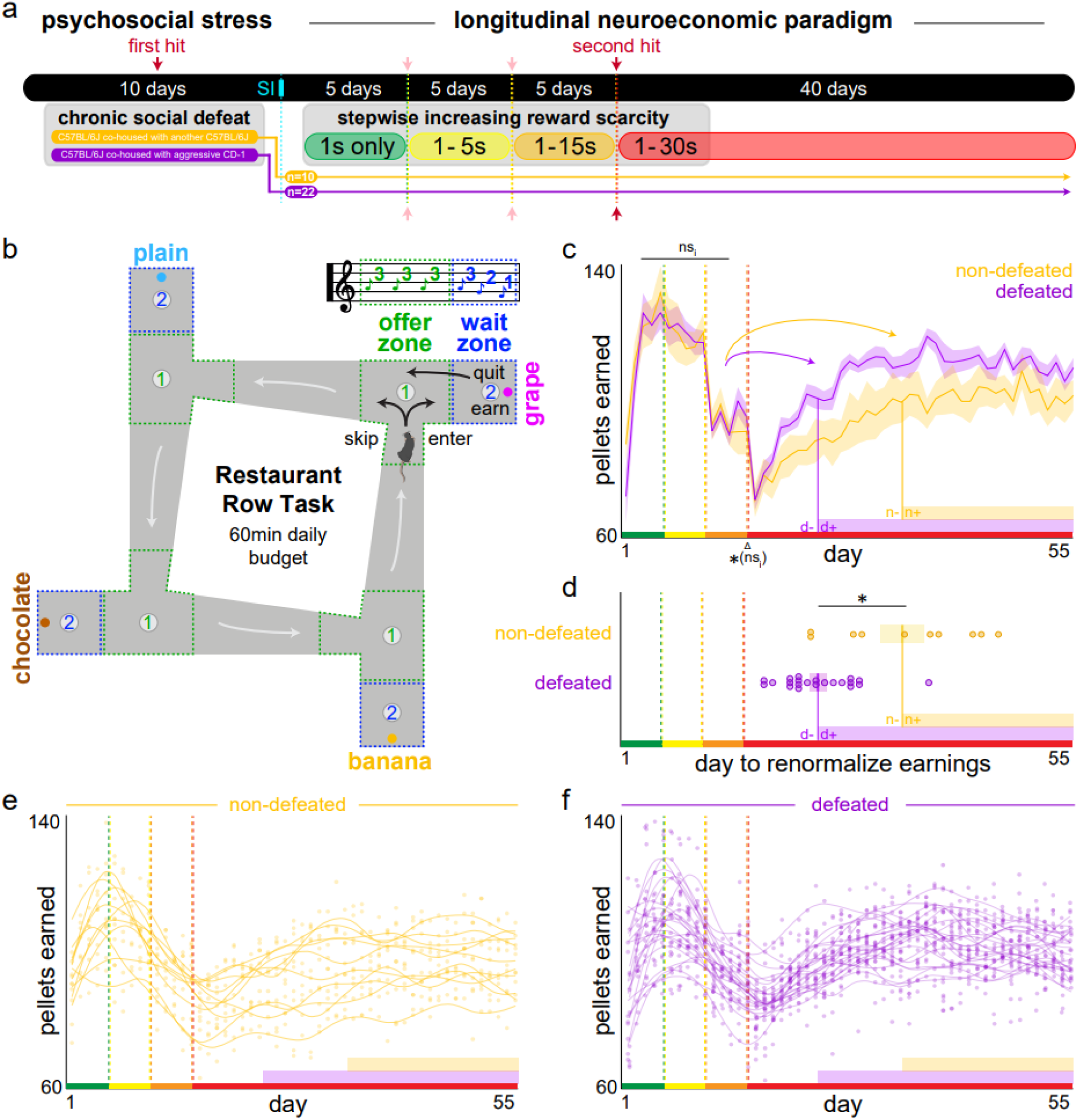
Novel two-hit social and economic stress model. (a) Experimental timeline. Mice were exposed to chronic social defeat stress (first hit, dark red arrow) followed by a brief social interaction (SI) screening assay before being tested longitudinally (55 d) in a spatial neuroeconomic decision-making paradigm with a changing value landscape that grew increasingly scarce with 5-d step (pink arrows). After 15 d, mice experienced the largest change in reward scarcity (economic crisis, second hit, dark red arrow) during which they abruptly transitioned into an environment where offers ranged from 1 to 30 s. (b) Restaurant Row task schematic. Mice were required to forage for their sole source of food while on a limited 60 min daily time budget for rewards of varying costs (delays) and flavors (four uniquely contextualized locations, or “restaurants”). Mice learned to run counterclockwise in a single direction encountering an offer zone in each restaurant in which a tone was presented on each trial whose pitch indicated the amount of time mice would have to wait for a reward should they choose to enter the wait zone, after which a pitch-signaled countdown would begin. (c) Average total number of daily pellets earned across the entire 55 d of testing, first in reward-rich environments where all offers were 1 s only (low pitch, green epoch), advancing stepwise into increasingly reward-scarce environments where offers could range from 1 to 5 s (yellow epoch), 1 to 15 s (orange epoch), or 1 to 30 s (red epoch). All mice suffered a drastic loss in food intake immediately upon transitioning into the 1 to 30 s reward-scarce environment (delta Δ symbol). Recovery back to near baseline levels of food intake (curved arrows, number of days to equate earnings with average yellow and orange epochs) for each group is indicated by solid, vertical drop lines where “-” indicates the window of testing before vs “+” after food intake was renormalized, summarized as number of days in (d) for individual animals. Purple and gold shaded bars along the x-axis indicate the “-” to “+” renormalization transition for each group throughout the remainder of figures as a point of reference. (e-f) Food earnings for individual mice across all 55 d of testing. Dots and smoothed lines represent individual animals. Shaded and x-y error bars ± 1 SEM. Not significant interaction (ns_i_).

### Neuroeconomic Decision-Making Task

We then characterized defeated and non-defeated mice longitudinally in Restaurant Row.^18^ Mice had a limited time period each day to forage for food by navigating a maze with four uniquely flavored and contextualized feeding sites, or “restaurants” (Fig. 1b). Each restaurant had a separate offer zone (OZ) and wait zone (WZ). Upon entry into the OZ, a tone sounded whose pitch indicated how long of a delay mice would have to wait in a cued countdown should they enter the WZ. Because animals were allotted only 1 hour to forage for their sole source of food for the day (full nutrition flavored BioServe 20 mg pellets), choices on this task were interdependent across days.

Thus, this task is economic in nature, requiring animals to budget their limited time effectively in a self-paced manner in order to earn a sufficient amount of food. Mice were tested across 55 consecutive days where strategies developed longitudinally as animals learned the structure of the task and the changing economic landscape. During the first 5 days of testing (block 1, green epoch), all trials consisted of 1 s offers only. During the next 5 days of testing (block 2, yellow epoch), offers ranged from 1 to 5 s randomly selected from a uniform distribution. Block 3 (5 days, orange epoch) consisted of a 1 to 15 s range. The fourth and final block (red epoch) consisted of offers ranging from 1 to 30 s and continued for 40 days. This block-wise protocol allowed us to examine how mice initially shaped their behavior as they learned the basic structure of the task as well as how they adjusted their foraging strategies across long timescales (days, weeks, and months) as they transitioned from reward-rich to reward-scarce environments in a stepwise manner. Rewards were delivered using a custom-built 3D-printed automated pellet dispenser that was triggered by a computer running the task programmed in ANY-Maze (Stoelting), run under dim lighting.

### Statistical analyses

Data were processed in Matlab with statistical analyses in JMP Pro 16. All data are expressed as mean ± 1 standard error. Statistical significance was assessed using student’s t tests and one-way, two-way, and repeated measures ANOVAs. Correlations were reported using Pearson correlation r coefficients. Akaike information criterion were used to compare model fits between linear versus cubic functions. No data were excluded as outliers.

## Results

### An economic crisis unmasks different foraging patterns in mice with a history of social stress

#### Food intake renormalization

Having a history of chronic social defeat stress had no immediately observable effect on task acquisition, feeding behavior, or body weight during early testing in relatively reward-rich environments in Restaurant Row (blocks 1-3, green-yellow-orange epochs, Fig. 1c, no interaction across the first 15 d: *F*_1,14_=1.802, *p*=0.180, Fig. S2). Each 5-d block increase in reward scarcity was marked by an abrupt transition into an environment with a larger offer range. Block four (red epoch) marked a critical point when mice experienced the greatest increase in offer range, up to 30 s, and experienced an immediately sharp, drastic loss in food intake to extremely low levels of earnings (with a concurrent drop in body weight, observed equally among defeated and non-defeated mice (Fig. 1c, main effect across the reward-scarce transition: *F*_1,1_=24.714, *p*<0.0001; no interaction by group: *F*_1,31_=0.227, *p*=0.635, Fig. S2). Interestingly, differences between defeated and non-defeated mice in total pellets earned by the end of each session were only observable after mice transitioned to the fourth block of testing. Over the subsequent weeks, mice recovered overall food intake back to levels near those observed in reward-rich environments. Surprisingly, mice with a prior history of social defeat stress renormalized food procurement in fewer days compared to non-defeated mice (Fig. 1c-f, *F*_1,31_=20.046, *p*<0.0001). These data show that an economic challenge is capable of extracting differences in foraging even after delayed timepoints in mice with relatively distant stress histories.

#### Energy expenditure and simple choice breakdown reveal multiple stages of a food-crisis response-profile

During testing in reward-rich environments, social defeat had no effect on overt locomotor behavior as all mice were able to learn to run the same number of laps in the correct direction (Fig. 2a, no interaction across the first 15 d: *F*_1,14_=0.454, *p*=0.501). Following the transition into a reward-scarce environment, there was a gradual and sustained increase in laps. Surprisingly, this effect was larger in defeated mice compared to non-defeated mice (Fig. 2a, *F*_1,31_=94.099, *p*<0.0001). The rise in laps was approximately in synch with the food renormalization window, respectively for each group of mice (recall “d-” and “n-” in Fig. 1c-d), where laps generally reached a plateau phase only after food intake renormalized for each group (recall “d+” and “n+” in Fig. 1c-d). This time course following the transition into the 1 to 30 s epoch, thus, can be divided into three distinct phases ([1] immediate, [2] delayed, and [3] long-term phases) when examining changes in simple choice outcomes (OZ: enter vs skip; WZ: earn vs quit). First ([1] immediate phase), there were no changes in OZ choice outcomes in either group of mice immediately following the transition into a reward-scarce environment (Fig. 2b, enters: *F*_1,31_=1.088, *p*=0.298; skips: *F*_1,31_=1.024, *p*=0.313). However, there was a sharp, immediate increase in WZ quit outcomes that was equivalent in both groups of mice (Fig. 2b, reward-scarce transition: *F*_1,31_=28.360, *p*<0.0001; no interaction with groups: *F*_1,1_=0.090, *p*=0.764). Second ([2] delayed phase), approximately during the food intake renormalization window for each group of mice (“d-” and “n-”), there was a gradual increase in number of enters, skips, and quits that was greater for defeated mice (Fig. 2b, three-way interaction between group and choice type [enter, skip, quit] across the renormalization window following the reward-scarce transition: *F*_2,3_=6.452, *p*<0.01). Third ([3] long-term phase), following food intake renormalization (“d+” and “n+”) and after laps reached a plateau, choice profiles surprisingly continued to change despite no obvious additional gain in food intake or decrease in energy expenditure (laps run). Most notably, skip outcomes continued to rise decoupled from any further changes in laps; and, unlike before during phase [2], was associated with a concomitant decrease in enters and quits. This third-phase effect was greater in defeated animals, and when normalized to laps, was disproportionally more pronounced than non-defeated mice (Fig. 2b-c, interaction of choices normalized to laps between zone and group during the last 2 weeks of testing: *F*_1,1_=57.717, *p*<0.001). These data highlight how an abrupt transition into a reward-scarce environment, which precipitates low yield in food intake, can reveal an immediate clash between behavior-environment interactions and how resultant poor performance can consequently drive multi-stage changes in foraging profiles across distinct timescales that will become more apparent with finer neuroeconomic analyses described below.

**Figure 2.**
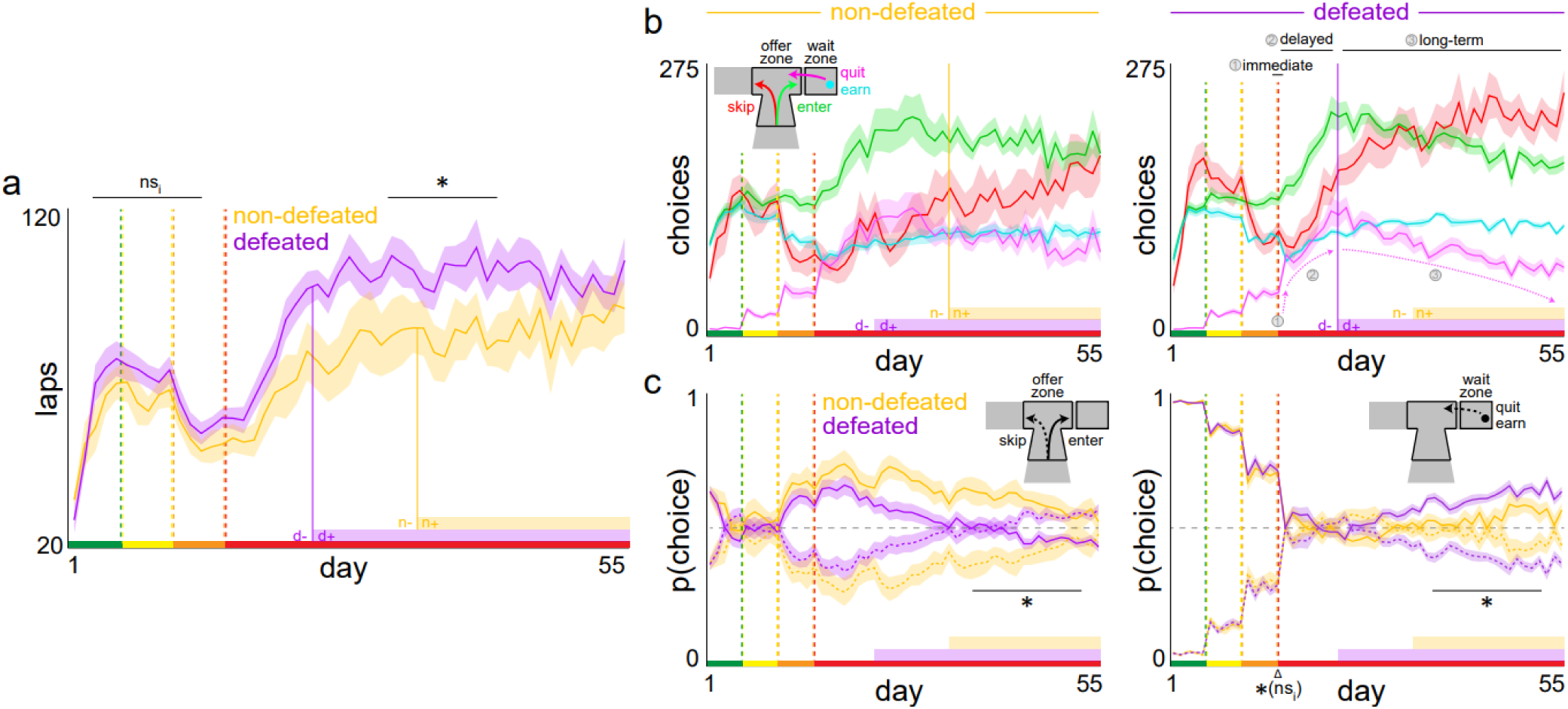
Energy expenditure and simple choice breakdown reveal multiple stages of a food-crisis response-profile. (a) Average number of laps ran in the correct direction. Note the “-” and “+” annotations along the purple (defeated, “d”) and gold (non-defeated, “n”) shaded bars along the x-axis demarcate food intake renormalization windows separated by vertical solid purple and gold lines as defined in Fig. 1c-d. (b) Average number of enter or skip choices made when in the offer zone as well as quit or earn choices in the wait zone for non-defeated and defeated mice. In addition to the “-” and “+” food intake renormalization window annotations, note additional annotations numbered [1] immediate phase (day 15 to 16 transition to 1 to 30 s offers, red epoch), [2] delayed phase (during the “-” renormalization window for each group of mice), and [3] long-term phase (after the “+” renormalization window for each group of mice). The magenta dashed arrow markings use quit behavior for defeated mice as an example to summarize and illustrate distinct changes in quit behavior associated with these three distinct phases (e.g., [1] immediate sharp rise, [2] delayed gradual rise, and [3] long-term gradual decline in quitting behavior). Note these changes in quits during these phases while considering changes in food intake, laps run, and enter vs skip choices. (c) The probability of making a choice in the offer zone relative to total number of offers or a choice in the wait zone relative to total number of enter decisions. Note that p(skip) and p(quit) here [dashed lines] are 1 minus p(enter) and p(earn) [solid lines], respectively, reflected about the 0.5 chance line (horizontal dashed gray line). Shaded error bars ± 1 SEM. Change relative to the transition into the 1 to 30 s reward-scarce environment (delta Δ symbol). Not significant interaction (ns_i_).

#### Redistribution of the limited time budget

In order to better grasp how mice responded to a decrease in reward availability upon transitioning into a reward-scarce environment, we next characterized how mice allocated their limited time budget to every action measured on this task. All behaviors on this task fall into one of 6 action-selection processes: time spent consuming rewards, waiting before earning, waiting before quitting, deciding to skip, deciding to enter, and traveling between restaurants (listed in order from greatest to least proportion of total session time allocated in experienced mice, Fig. 3a). We measured each metric as a percentage of total session time (Fig. 3a-b) as well as average time spent engaged in a single bout (Fig. 3c). We found a complex restructuring of time engaged in these various behaviors across the entire experiment in response to economic stress (Fig. S3). Mice with a history of social defeat differed from non-defeated mice in these time-related metrics only after the transition into a reward-scarce environment. History of social defeat stress influenced changes not only in the proportion of total session time spent engaged in a particular behavior following economic stress, which is related to alterations in event frequency as displayed in Fig. 2, but also in average time spent engaged in a single behavioral episode. Of note, we found social defeat-related differences in the two reward-rejecting action-selection processes – skipping and quitting behavior – in both frequency and average duration compared to non-defeated mice. We found that over long timescales after the transition into a reward-scarce environment, compared to non-defeated mice, defeated mice displayed a greater increase in proportion of total session time spent skipping while conversely spending a smaller proportion of total session time spent quitting. Furthermore, defeated mice learned to make these decisions faster than non-defeated mice. Without differences in proportion of session time spent entering between groups of mice, these data indicate defeated mice undergo a more robust change in decision-making strategies rooted in a shift in reward-rejecting policies between the OZ and WZ. Because opportunity costs carry more weight in a reward-scarce environment, change-of-mind decisions in the WZ and skip-first decisions in the OZ, which are thought to employ computationally distinct valuation algorithms^21,22^, may be differentially perturbed in defeated mice when considering foraging elsewhere for food.

**Figure 3.**
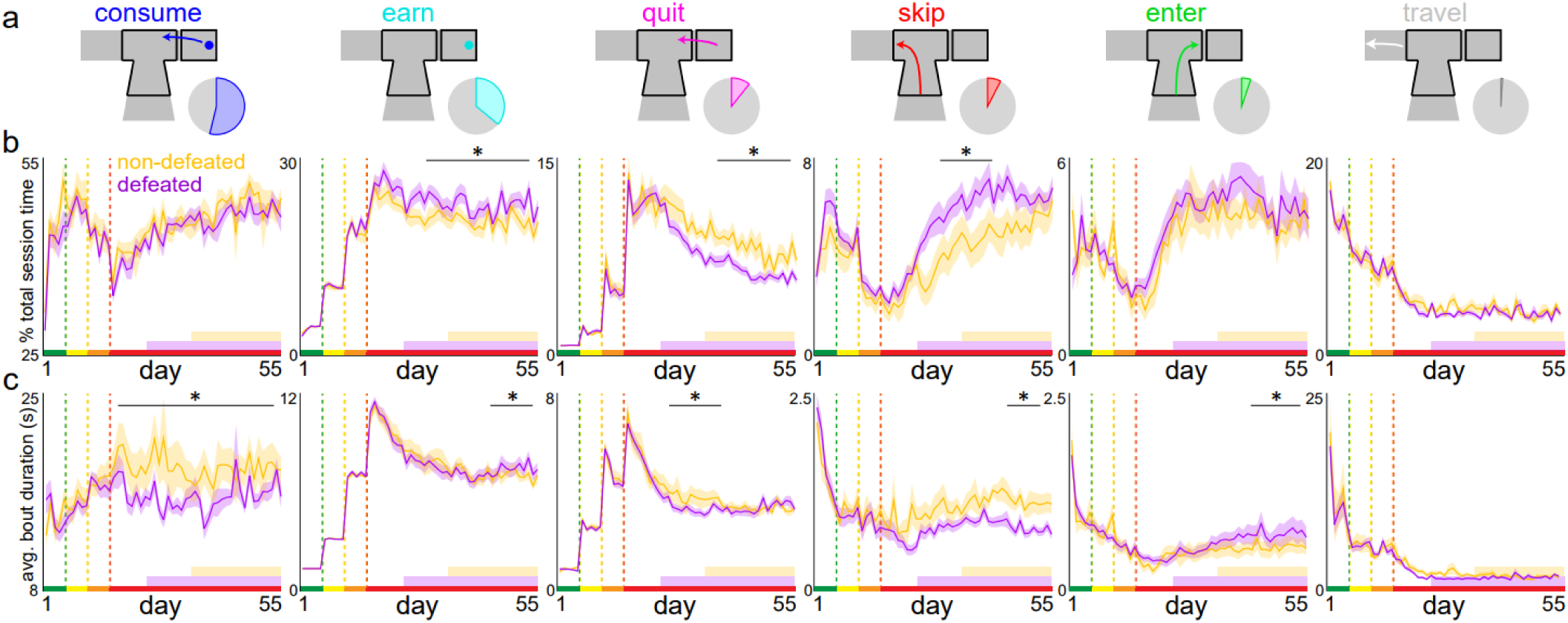
Redistribution of the limited time budget. (a) Schematics of all possible and mutually exclusive behaviors in which mice could be engaged on the Restaurant Row task sorted by total proportion of time allocated represented in the accompanying pie charts (average of the final 5 d of testing). (b) Percent of total daily session’s time budget allocated to each behavior across all 55 d of testing for defeated and non-defeated mice. (c) Average duration of a bout of each behavior across testing. See Fig. S3 for summary statistics. Shaded error bars ± 1 SEM.

#### Learning occurs across multiple, economically distinct, valuation algorithms

To quantify the economic nature of the types of decisions being made, we next analyzed how cost and time-related elements interacted to drive choice behavior. This is particularly important to understand as mice appeared to counterintuitively increase energy expenditure in a reward-scarce environment by running more laps, more so in defeated mice. First, we calculated thresholds of willingness to wait in each zone by fitting a heaviside step regression to choice outcome as a function of cued offer cost on each day (Fig. 4a). OZ thresholds reflect willingness to enter initially presented offers, whereas WZ thresholds reflect willingness to remain committed to an ongoing investment upon entering. A large discrepancy between OZ and WZ thresholds immediately emerged following the transition into a reward-scarce environment, where OZ thresholds drastically increased to nearly 30 s (Fig. 4b, main effect across the reward-scarce transition: *F*_1,1_=52.836, *p*<0.0001; no interaction by group: *F*_1,31_=0.001, *p*=0.970). This means that animals indiscriminately entered virtually all offers in the OZ regardless of cost (Fig. S4a,c), a policy they previously adhered to without consequence in reward-rich environments. The large discrepancy with WZ thresholds describes that mice accepted a large proportion of offers they were not actually willing to wait for once in the WZ, hence precipitating the sharp increase in quit outcomes (recall Fig. 2b-c, immediate phase). This can explain why mice suffer an immediate loss in food intake – a significant portion of their limited time budget was “wasted” to quitting behavior. This also explains why during the delayed phase (during the “-” renormalization window), as laps increased, entering, skipping, and quitting behavior gradually increased over the subsequent weeks because OZ economic policies remained relatively fixed. This response profile during the delayed phase timepoint, although at the expense of spending energy by running more laps, allowed animals to sample the new range of offers in a reward-scarce environment with greater frequency. This captures the development of a rigorous foraging strategy that essentially ignores offer cost in the OZ during the delayed phase but nonetheless is sufficient to renormalize food intake following an economic crisis in the short term.

**Figure 4.**
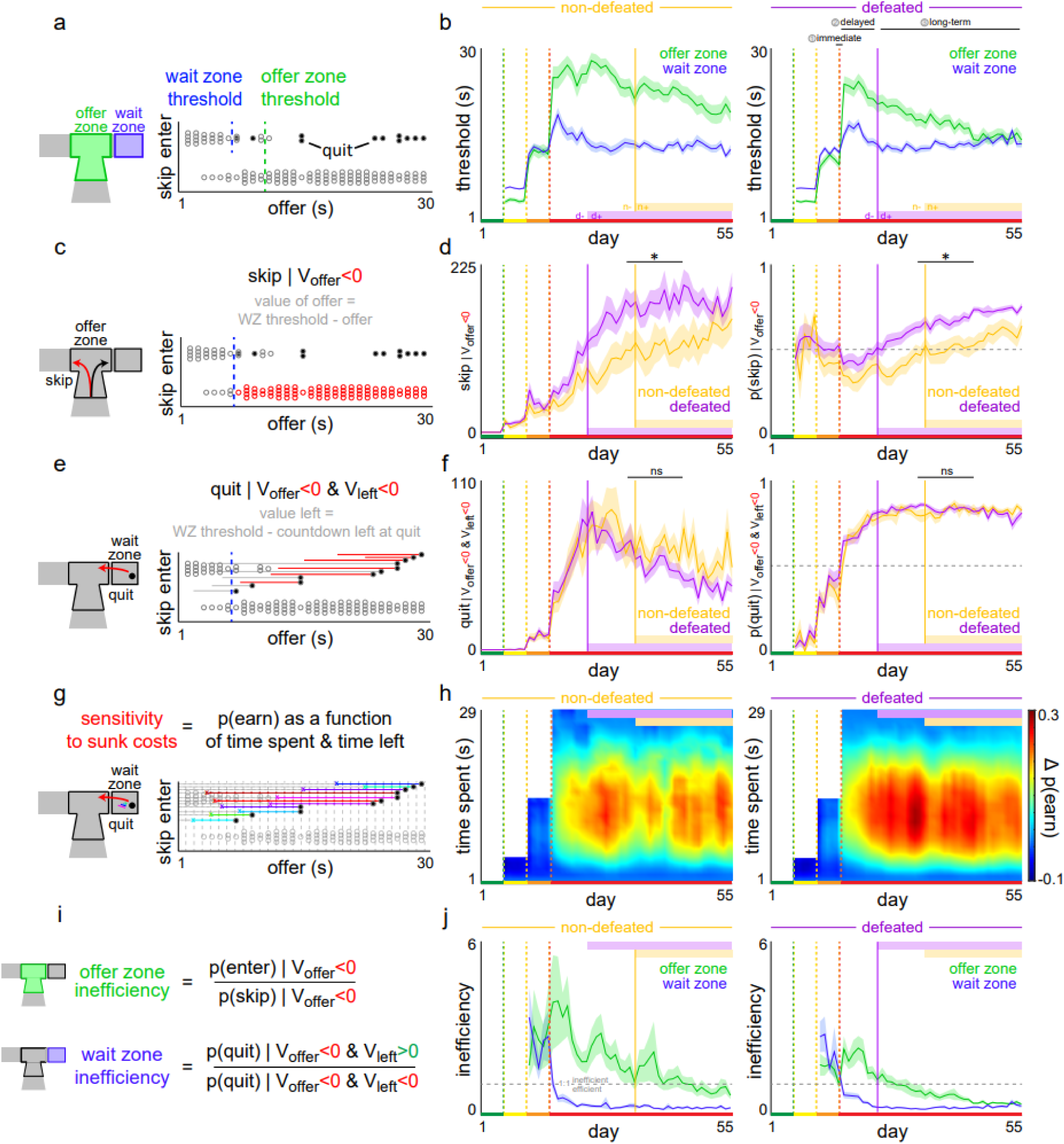
Learning occurs across multiple, economically distinct valuation algorithms. (a) Thresholds of willingness to wait are determined by fitting a heaviside step regression to choice outcome (enter vs skip in the offer zone or earn vs quit in the wait zone) as a function of cued offer cost (Fig. S4). Vertical dashed lines reflect example thresholds in the offer zone (green) or wait zone (blue) for a single restaurant from a single mouse in a single session. Individual trials in this example are represented by dots, with quits as filled black dots. (b) Average offer zone and wait zone thresholds across all 55 d of testing for non-defeated and defeated mice. (c) Since wait zone (WZ) thresholds are generally stable across testing for both groups of mice, offer value can be calculated relative to thresholds (gray text). (d) The average number of skip decisions and the probability of skipping given the offer is above threshold, where offer value (V_offer_) <0. (e) For quit decisions, the value left at the moment of quitting can also be calculated relative to thresholds (gray text). (f) The average number of quit decisions and the probability of quitting given the offer is above threshold, where V_offer_<0, and the time left at the moment of quitting is still above threshold, where value left (V_left_) <0. (g) Sensitivity to sunk costs can be calculated by comparing the probability of earning a reward in the wait zone as a function time already spent and time left in the countdown (see Fig. S5 for a visual explanation and quantification steps). (h) A heatmap of the change in p(earn) in the wait zone as a function of time spent (sunk cost sensitivity) independent of time left showing an escalation of commitment to waiting that is stronger the more time has already been waited. (i-j) Offer zone and wait zone inefficiency can be summarized by taking the ratio of probabilities of making an economically disadvantageous decision relative to the advantageous choice in the offer zone given V_offer_<0 or in the wait zone given V_offer_<0 depending on V_left_. Horizontal dashed gray line indicates a 1:1 ratio where policies become more efficient and fall below this line across testing. Shaded error bars ± 1 SEM. Not significant interaction (ns_i_).

To quantify these economic changes more precisely, we converted offers into value terms by normalizing trials to an individual’s WZ threshold for each flavor, which is generally stable across testing. That is, if a mouse’s threshold on a given day and in a given restaurant is 15 s and an offer on a given trial is 20 s, value_offer_= threshold – offer, or -5 in this example trial, reflecting an offer that is a “bad” deal and should be rejected (Fig. 4c). This allows individual differences in willingness to wait across mice or between flavors to become normalized into comparable value terms. With this approach, we found that skipping negatively valued offers did not begin to increase until late in testing and was more robust in defeated mice (Fig. 4d, across days: *F*_1,39_=232.307, *p*<0.0001; between groups: *F*_1,31_=148.619, *p*<0.0001). This change underlies the gradual decrease in OZ thresholds observed during the long-term phase as defeated mice more rapidly learned to make cost-informed choices in the OZ by discriminating tones (Fig. 4b, across days: *F*_1,39_=165.084, *p*<0.0001; between groups: *F*_1,31_=168.530, *p*<0.0001, Fig. S4b,d). We applied a similar analysis to quit choices and included a term that captured the value remaining in the countdown at the moment of quitting. That is, if a mouse’s threshold on a given day and in a given restaurant is 15 s and an offer on a given trial is 20 s (i.e., a “bad” deal) but the animal quit after waiting only 3 s, with 17 s remaining in the countdown, value_left_ = threshold – time left, or -2 in this example trial, reflecting an accepted offer that was quit in an economically advantageous manner fast enough such that the deal remaining was “still bad” (Fig. 4e). This implies that if mice accepted bad deals in the offer zone, they were largely able to quit these trials quickly, correcting OZ mistakes. We found that these economically efficient quits gradually increased during the delayed phase after transitioning into a reward-scarce environment, in parallel with a decreased latency to quit (recall Fig. 3c), but with no differences between defeated and non-defeated mice (Fig. 4f, across days: *F*_1,39_=233.364, *p*<0.0001; no effect between groups: *F*_1,31_=3.310, *p*=0.069). This early rigorous foraging strategy typified by a high rate of change-of-mind decisions was sufficient to renormalize food intake during the delayed phase. At later long-term timepoints after food intake had renormalized, quitting behavior decreased as mice learned to effectively trade an enter-then-quit strategy for a skip-first strategy in the OZ. The economic efficiency of these OZ and WZ choices are summarized in Fig. 4i-j, which highlights separate learning timescales of each decision process.

Focusing more on quitting behavior, the notion of time spent while waiting during the countdown for a reward is an intriguing concept and a perplexing economic dilemma. Mice must make a time-sensitive decision to quit an ongoing investment from a limited budget while progressing closer toward earning a reward, re-evaluating whether or not to change one’s mind. This scenario captures the economic phenomenon known as the sunk cost bias, where individuals tend to overvalue and escalate commitment of an ongoing endeavor as a function of irrecoverable losses, with stronger effects growing with larger time investments.^23^ We developed a dynamic analysis capable of extracting sensitivity to sunk costs in the form of time by calculating the probability of staying versus quitting in the WZ as a function of both time spent and time left remaining in the countdown (Fig. 4g).^24,25^ Sensitivity to sunk costs can be calculated as a change in the probability of earning [Δp(earn)] depending on how much time was spent relative to conditions in which mice waited 0 s (see Fig. S5a-d for a visual explanation). We found that all mice become sensitive to sunk costs only in reward-scarce and not in reward-rich environments (Fig. 4h, yellow & orange epochs: sign test Δp(earn) > 0: *t*=-21.192, *p*=0.999; red-epoch: sign test Δp(earn) > 0: *t*=+112.005, *p*<0.0001). Furthermore, we found that defeated mice display enhanced sensitivity to sunk costs and, in particular, demonstrated an increased rate at which sensitivity to sunk costs developed across days of testing (Fig. 4h, Fig. S5e-f, *F*_28,39_=17.929, *p*<0.0001). These data indicate that multiple valuation algorithms change in response to an economic crisis and to different degrees depending on whether or not animals have a prior history of social stress.

### Strategies that optimize food security can also separately contribute to subjective value

Given that economic decision policies continue to change through the end of this experiment despite no obvious additional gain in food intake or decrease in energy expenditure, we next explored how individual differences in subjective value may play a role in the additional learning that may be guiding these further shifts in strategy. Note that the Restaurant Row task employs the use of different flavors in each restaurant as a way, separate from cost, to manipulate individual differences in subjective value (as opposed to varying pellet number, for example, which would introduce more complexities and added costs due to increased handling time to consume different numbers of pellets). Daily preferences here were measured by simply adding the total number of pellets earned in each restaurant at the end of each session and ranking flavors from least to most preferred. While each mouse has a preferred flavor, there was no group difference across mice in these preferences. During testing in rewardrich environments, social defeat had no effect on revealed flavor preferences (Fig. 5a-b, across green-yellow-orange epochs: no interaction between group and earns by flavor: *F*_1,31_=0.411, *p*=0.745; no effect of group on standard deviation of flavor earnings: *F*_1,31_=1.969, *p*=0.161). By contrast, the transition into a reward-scarce environment had the greatest impact on earnings for most preferred flavors, with more robust recovery for these flavors in defeated mice compared to non-defeated mice (Fig. 5a, most preferred flavor: *F*_1,31_=180.749, *p*<0.0001; least preferred flavor: *F*_1,31_=2.177, *p*=0.140).

**Figure 5.**
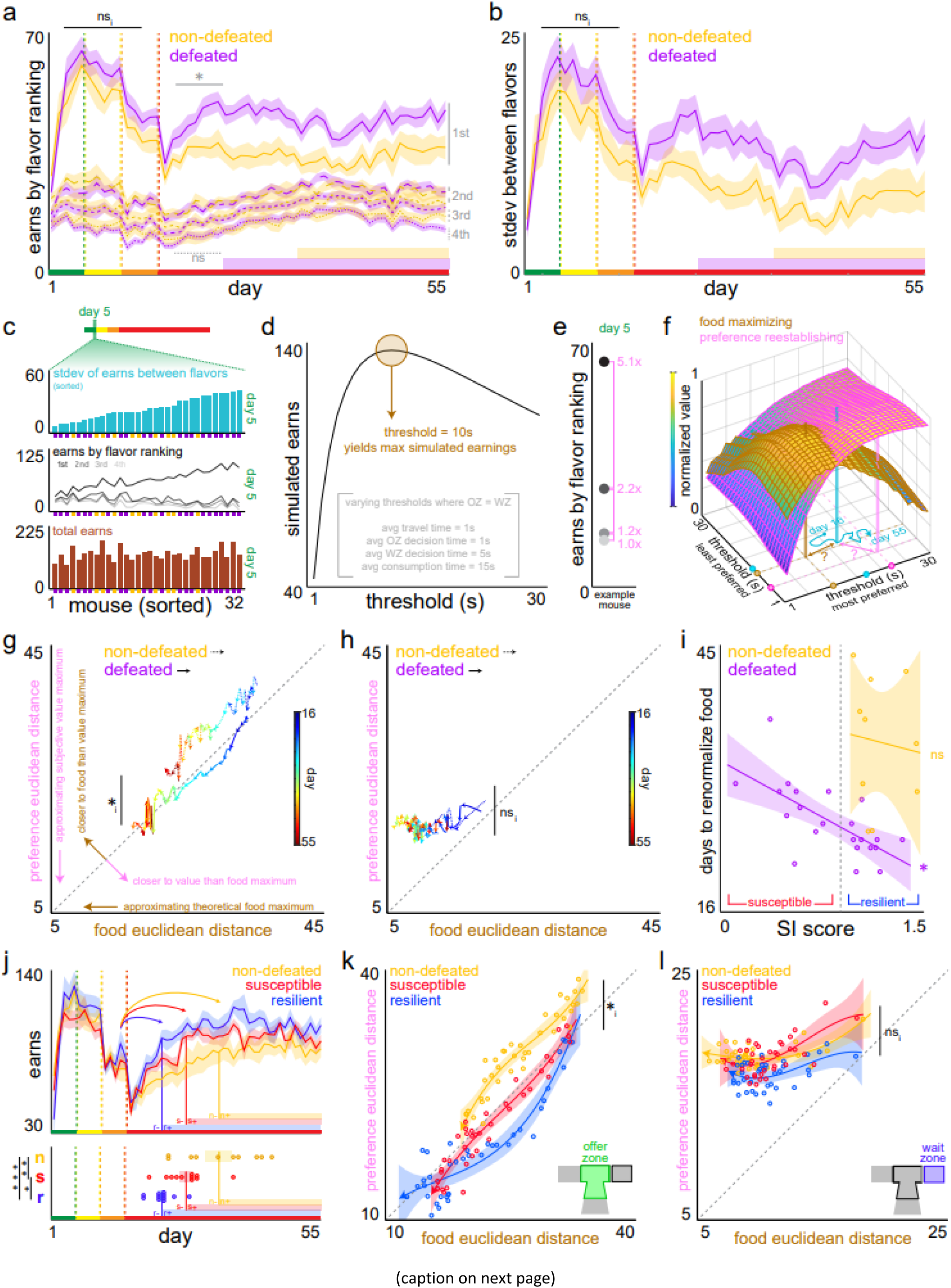
Decision policies that optimize food security are shared with but can also independently guide strategies that separately promote subjective value. (a) Average number of pellets earned in each restaurant ranked from most to least preferred across 55 d of testing. (b) Average standard deviation of earns among the four flavors. (c) Display of individual mouse behavior (standard deviation of earns across the four flavors, total number of earns by ranked flavors, and total number of overall earns), extracted only from day 5 of testing, the final day of the green epoch where all offers were 1 s only. These day 5 data were sorted by the standard deviation of earnings across flavors, from similar to dissimilar flavor palettes. Purple and gold squares along the x-axis denote defeated or non-defeated membership. (d) Computer Restaurant Row simulation of total number of pellets earned. The ideal threshold required to obtain the theoretical maximum number of pellets when ignoring flavor preferences was determined to be 10 s. (e) Example session: number of pellets earned across flavors ranked most (darker) to least preferred from a single mouse extracted from day 5. Pink inset text indicate the relative ratio of earns for this mouse on day 5 between the flavor rankings (i.e., this mouse had a 5.1 : 2.2 : 1.2 : 1 parts recipe across flavors for idealized subjective value when all offers were 1 s only). (f) Two intersecting planes of decision policies that yield varying amounts of theoretical value either for maximal food intake (as determined by computer simulations, brown plane) or subjective value (as determined by day 5 preferences on a mouse-by-mouse basis, pink plane) when in a reward-scarce environment (1 to 30 s offers). Here, only two decision policy dimensions (least preferred and most preferred restaurants) of the four dimensions (all four ranked restaurants) are displayed. Both planes are normalized to each’s min and max values for the purpose of plotting both in the same space while preserving the threshold coordinate locations that achieve either theoretical maximum (brown [location always fixed at threshold coordinates of 10 s] or pink beacons [location of discovered threshold coordinated vary from mouse-to-mouse]). Actual observed daily decision policies represented by the cyan beacon wander throughout this space from day 16 to 55 in a reward-scarce environment. Trajectories projected to the floor of this display trace out individual mouse decision policy paths. Example coordinates of brown, pink, and cyan beacons illustrated as dots on the x and y axes. Euclidean distance from cyan coordinates to either brown or pink coordinates were calculated (question mark symbols). (g-h) Scatter plot of Euclidean distances from observed decision policies in the offer zone (g) or wait zone (h) to either food or preference theoretical maximum across testing in a reward-scarce environment (color-coded arrowed line stepping through days, non-defeated mice in dashed arrows, defeated mice in solid arrows). See Fig. S6 for more comprehensive and summary data. (g-h) intended to highlight progression across days while (j-k) below highlight trajectory shape. (i) Scatterplot of food intake renormalization rates and social interaction (SI) score. Vertical dashed gray line indicates an SI score of 1. Dots represent individual mice. (j) Food intake renormalization data from Fig. 1c-d replotted splitting defeated mice by resilient and susceptible subgroups (as defined by SI score >1 or <1 respectively). (k-l) Decision policy trajectories across testing replotted with resilient and susceptible subgroups. Pointed arrowhead indicate overall direction across days of testing as shown in (g-h). Dots represent group mean on each day. Diagonal dashed gray lines in (g-h) and (k-l) indicate a slope of 1. Shaded and x-y error bars ± 1 SEM or 95% CI of curve fits. Not significant interaction (ns_i_).

The fifth day of the first epoch of testing (day 5, 1 s only offers, green epoch) presents a useful timepoint when data can be harnessed to examine how decision-making policies that develop across the remainder of the experiment fare in reapproximating the distribution of the relative number of pellets earned among flavors compared to day 5. Examining data in this way allows us to ask how animals might prioritize subjective value across testing. Compared to when costs were at an all-time low after mice have already learned the basic structure of the task and with stabilized behavior on day 5, we asked how the distribution of relative flavor preferences related to changing decision policies once in a reward-scarce environment later in testing. On day 5, when rewards were essentially “free,” mice displayed a broad range of individual differences in flavor preferences unrelated to total earns or defeat history (Fig. 5c). Next, using a computer simulation of the Restaurant Row task, holding several behavioral measures constant, we calculated the maximal number of pellets that could be theoretically earned as a function of threshold if a mouse had no flavor preferences and equivalent OZ and WZ policies (Fig. 5d). We found that a threshold of 10 s is the optimal amount of time one should be willing to accept and wait in all restaurants in a reward-scarce environment (when offers range from 1 to 30 s) if the sole objective is to purely maximize food intake (Fig. 5d). With this in mind, we next aimed to capture a separate theoretical maximum of decision thresholds, not for objective food acquisition, but instead for subjective value of food that is unique to an individual mouse. To do so, we performed another computer simulation of the Restaurant Row task where thresholds for flavors ranked from least to most preferred were modeled separately and independently. We extracted the ratio of relative number of pellets earned in each restaurant as defined on day 5 per mouse (Fig. 5e). This allowed us to define a subjective “preference” value by multiplying reward outcomes with mouse-specific flavor preferences (defined by the day 5 ratio of flavor earnings). We can then visualize both the objective food intake and subjective “preference” value as a function of thresholds in each restaurant (3D surfaces displayed for the most preferred and least preferred restaurants in Fig. 5f). Thus, we generated different combinations of decision-making thresholds across restaurants in a reward-scarce environment that map how much food in theory could be earned (food maximizing) versus how much total mouse-specific value could be obtained (preference reestablishing, Fig. 5f). From this, we identified the coordinates of decision thresholds across restaurants in a reward-scarce environment that achieve (i) maximal theoretical food (i.e., always 10 s in all 4 restaurants) or (ii) maximal theoretical subjective value (a set of thresholds in each restaurant unique to each mouse). Finally, mouse-by-mouse, we plotted the actual set of thresholds in each restaurant on each day in order to observe the trajectory through these planes across days of testing in a reward-scarce environment (cyan trace projected to the floor of the visualization in Fig. 5f).

Using this analysis, we asked how the decision policies of defeated and non-defeated mice prioritize maximizing food intake versus subjective values across testing following an economic challenge. We calculated the Euclidean distance between the observed thresholds and either optimal food-intake thresholds or optimal subjective value thresholds on each day in both the OZ and WZ (Fig. 5f). We found a significant difference in decision-making trajectories through these planes across testing in a reward-scarce environment between defeated and non-defeated mice depending on the type of choice being made. In the OZ, all mice decreased Euclidean distances to theoretical maxima on both the food intake as well as the preference reestablishing planes across days of testing (Fig. 5g, main effect of day on distance across groups: *F*_1,39_=438.753, *p*<0.0001, Fig. S6a-b,e,g). However, defeated mice more robustly drove both distances closer to absolute zero across days of testing (main effect of group: *F*_1,31_=268.423, *p*<0.0001; interaction between group and distance type: *F*_1,31_=38.699, *p*<0.0001, Fig. S6a-b,e,g). Groups of mice did not differ in distances between observed thresholds and either theoretical maxima early in testing in a reward-scarce environment (Fig. S6g). Groups of mice also did not differ in distances between theoretical food maximum and theoretical subjective “preference” value maximum (Fig. S6h). By subtracting Euclidean distances in OZ thresholds between both theoretical maxima to measure relative distances, we found that defeated mice decision policies were closer to theoretical subjective value maxima than to theoretical food maxima compared to non-defeated mice (Fig. 5g, Fig. S6c). Interestingly, in the WZ, decision-making trajectories traversed these planes in a manner that decreased the Euclidean distance only to food intake policies, with virtually no contribution to reestablishing subjective value preferences and with no differences between groups of mice (Fig. 5h, interaction between day and plane: *F*_1,39_=35.183, *p*<0.0001; no main effect of group: *F*_1,31_=1.163, *p*=0.281, Fig. S6a-b,d,f,g). These data indicate that subtle changes in OZ and WZ thresholds can contribute to hidden aspects of food- or value-related gains that might not be readily appreciable if not considering cross-restaurant policies and within-subject behavior relative to a reward-rich environment.

Finally, in order to link unique social defeat stress response profiles to economic stress response profiles, we correlated social interaction (SI) scores with food intake renormalization rates and found that defeated mice with higher SI scores required fewer days to renormalize following the transition into a reward-scarce environment (Fig. 5i, defeated: *F*_1,20_=16.43, *p*<0.001; non-defeated: *F*_1,8_=0.08, *p*=0.787, Fig. S1b). High SI scores following social defeat (approach behaviors) have traditionally been linked to resilient stress-response phenotypes on numerous simple behavioral screens, whereas mice with low SI scores (avoidance behaviors) have traditionally been labeled as stress-susceptible and exhibit depressive-like traits in other assays.^17^ We found that resilient mice displayed the fastest food intake renormalization rates following the transition into a reward-scarce environment compared to susceptible and non-defeated mice (Fig. 5j, *F*_1,2_=13.985, *p*<0.001; *t*_non-defeated v resilient_=5.28, *p*<0.0001; *t*_non-defeated v susceptible_=3.06, *p*<0.01; *t*_resilient v susceptible_=-2.27, *p*<0.05). Extending our decision-making trajectory analysis to these subgroups of mice, we found a striking bifurcation in Euclidean distances only in the OZ but not WZ (Fig. 5j-l). While all groups followed generally similar decision trajectories in the WZ to decrease distance only to the theoretical maxima for food intake (Fig. 5l, main effect of day on food plane: *F*_1,39_=53.866, *p*<0.0001; no main effect of day on subjective value plain: *F*_1,39_=0.889, *p*=0.346; no interaction between groups and distance type: *F*_1,1_=0.928, *p*=0.395), resilient mice uniquely steered OZ trajectories more closely toward decision policies that decreased the Euclidean distance to preference reestablishing maxima while simultaneously driving the Euclidean distance to food maxima down (Fig. 5k, interaction between groups and distance type: *F*_1,1_=31.649, *p*<0.0001, food vs preference trajectory model curve fit secondary Akaike information criterion (AICc) weights: non-defeated [linear: 0.721 ; cubic: 0.279]; resilient [linear: 0.004; cubic:

0.996]; susceptible [linear: 0.878; cubic: 0.122]). Subgroups of mice did not differ in distances between theoretical food maximum and theoretical subjective “preference” value maximum (Fig. S6i). These data indicate that mice with a history of a resilient profile in response to a prior social stressor are more capable of recovering from an economic setback. Resilient mice accomplish this in a manner that not only ensures food security necessary for survival but also better prioritizes subjective value. Lastly, differences in decision-making policy trajectories appear to emerge only in certain types of choices (OZ but not WZ).

## Discussion

The present study employed a novel two-hit stress model in order to examine how the decision-making processes involved in recovering from an economic setback are altered in mice that have a history of prior social stress. We found that mice exposed to social defeat when tested longitudinally on a neuroeconomic task mounted a more robust behavioral response than non-defeated mice only after abruptly transitioning into a reward-scarce environment. We found that the magnitude and learning trajectory of the decision-making response to this economic challenge differed between mice that were resilient versus susceptible to the initial social stress, with resilient individuals optimizing the recovery of both food intake and subjective value.

Economic stress is widely considered one of the most pervasive and universal burdens to mental health in the human experience, made worse by the COVID-19 pandemic and current inflationary period.^16^ Economic crises, including job loss and recessions, are associated with increased use of mental health services as well as increased mortality and suicide rates.^26-29^ Financial strain can lead to impaired functioning, poor overall physical health, and mood disorders such as depression.^30-32^ Economic stress can also precipitate first-episode mental illness in at-risk or otherwise previously healthy individuals.^11^ Our knowledge of the neurobiological underpinnings of economic stress in psychiatric disorders is limited, however, several reports in the human literature have linked effects of recent financial but not other types of stressors to (i) polymorphisms in the serotonin transporter gene, (ii) longlasting connectivity changes in the default mode network associated with increased activity of the hypothalamic-pituitary-adrenal axis during social stress, and (iii) even lower remission rates during antidepressant treatment.^33-36^ Virtually no animal studies to date have characterized the effects of economic stress on behavior in a psychiatric disease model. In general, animal tasks that involve changing rules, contingencies, or effort demand indeed tax the individual in ways that pressure shifts in behavior toward strategies that may be less costly, as is commonly seen in traditional operant tasks with escalating work schedules or in reversal learning paradigms.^37-42^ However, rarely are rewards in such tasks critical for survival. Furthermore, few studies have examined how changes in task rules are necessary in order to extract group differences in animal models used for the study of stress-related disorders – differences that would otherwise go unobserved when task demand is low.^43-45^ Our novel two-hit model for studying economic stress, in mice with a prior history of social stress, challenges animals with an abruptly severe and striking change in reward availability in the environment without compensating for fixed budget constraints that, in a closed-economy system, have dire consequences and to which mice must adapt longitudinally in order to survive.

The question about whether or not the behavioral changes following the economic challenge are adaptive or maladaptive requires close examination. Defeated mice demonstrated a faster response profile when renormalizing food intake compared to non-defeated mice. This can be interpreted in several ways depending on whether or not this is considered an abnormal response. Ultimately, the direction of this change is arguably in the best interest of these animals and favorable from a survival standpoint. Stress responses classically follow an inverted U-shaped relationship where too little or too much stress exposure, as well as the degree of the individual’s response to stress, can be deleterious.^46^ Where along this inverted U-shape curve mice with or without a history of social defeat fall when experiencing economic stress is debatable. It is possible that non-defeated mice were, overall, under less pressure to perform following the economic challenge given no other prior history of stress and consequently their renormalization rates sufficed for them. On the other hand, if a slowed response is abnormal, non-defeated mice might be achieving weaker renormalization rates because, in the absence of a prior adverse experience, they had no prior challenges to overcome as did the defeated mice. This interpretation seems less favorable because behavioral changes after experiencing social defeat are often thought to reflect impaired function overall where repeated bouts of stress generally sensitize responses to future stress rather than promote stress tolerance, although mixed findings have been reported in the literature.^47-49^ Nonetheless, in this task, energy conservation and metabolic needs, which are known to be altered following social defeat, may play more complex roles in driving different changes between groups of mice.^17,50-54^ For instance, the enhanced sensitivity to sunk costs that we observed in defeated animals could reflect several changes: (i) animal learning theory models purport that sensitivity to sunk costs could be driven by statedependent valuation learning or within-trial contrast whereby ongoing energy expenditures and caloric deficits that grow with the passage of time enhance the value of food rewards; (ii) sensitivity to sunk costs are known to increase under conditions of uncertainty and in reward-scarce environments, as observed here, and although within a single trial may be economically disadvantageous, across trials may be time-saving and cache more rewards in the long-run.^55-58^ Both of these factors could be exaggerated in defeated mice, either via larger caloric deficits or with greater perceived uncertainty in a reward-scarce environment, driving an escalation of commitment during change-of-mind decisions in the WZ. In any event, the differences in multiple valuation algorithms that emerge across testing suggest that complex responses to economic stress diverge between mice with and without a history of social defeat, including both resilient and susceptible subgroups.

Stress resilience can be conceptualized as a preservation in homeostasis where resilient individuals could be equated to those who did not experience stress at all.^2-6,48^ Practically, this could hold merit as resilient individuals, for example after trauma, do not meet clinical criteria for psychiatric diagnoses and would otherwise fall into the healthy population.^1^ However, this viewpoint has begun to fall out of favor in recent years as newer behavioral and neurobiological evidence shows that the manifestation of resilience is an active process that is necessary in order to protect against poor outcomes for both physical and mental health.^59-61^ In this context, just the right amount of stress, or “eustress,” can be healthy and as needed drive adaptive coping mechanisms in individuals with intact systems or without predisposed vulnerabilities.^62,63^ Stress susceptibility on the other hand has been framed in several ways, including characterizing stress responses as either an actively dysfunctional and maladaptive process or an absence of resilient adaptations.^17,64^ We characterized mice as susceptible versus resilient along a continuum of social-stress-induced social avoidance as classically measured in the literature, a well-validated predictor of depressive-like phenotypes on numerous other behavioral screens.^17,65^ We discovered that resilient mice were able to renormalize food intake the fastest in response to the second, economic hit upon transitioning into a reward-scarce environment compared to susceptible and non-defeated mice. Although susceptible mice too renormalized faster than non-defeated mice, their intermediate profile suggests that susceptible mice may be mounting a partial or exhausted stress response following the second, economic hit. Furthermore, the different decision policy trajectories among non-defeated, resilient, and susceptible mice highlight how all three groups weigh the balance between food security and subjective value in unique ways throughout their journey during the recovery process in a reward-scarce environment. In fact, the susceptible group’s shift in OZ trajectories away from theoretical value maxima relative to resilient mice may reflect a more complex and subtle take on anhedonia often reported in animals with a stress-susceptible phenotype.^17^ Although frequently studied in addiction, little is known about how elasticity of demand curves change in mood disorders, particularly when comparing essential goods to luxury items.^66-70^ Our data show that resilient individuals can better promote subjective value without sacrificing gains in food intake, although it is still intriguing that we find such differences in decision policies only in the OZ and not WZ.

Choices made between the OZ and WZ are thought to reflect fundamentally distinct decision-making algorithms.^18,21,22,24,71,72^ Mice learn to discriminate tones and make economically advantageous decisions in the OZ late in testing, generally after food intake has renormalized. This suggests that mice may be using cost-related information in ways that are not necessarily entirely in service of maximizing food intake.^18,20^ Previous reports with hippocampal tetrode recordings in rodents and functional magnetic resonance imaging of the default mode network in humans tested on translated versions of the Restaurant Row task found signatures of an ongoing, prospective deliberative decision-making process where individuals were more likely to represent future competing options before making a choice.^71,73-75^ The learning that takes place in the trajectory of OZ decision policies of resilient mice were more likely to factor in subjective value while simultaneously maximizing food consumption. It is possible that part of the prospective deliberative processes engaged by resilient animals while making cost-informed choices involves contemplating flavor more earnestly, ultimately contributing to more gains in subjective value over their susceptible and non-defeated counterparts. Change-of-mind decision policies in the WZ on the other hand, which seem to entirely promote food intake with no differences between groups of mice, have been shown to be uniquely affected by manipulations of the medial prefrontal cortex without altering OZ choices.^21,74,76^ These data suggest that physically distinct circuits, which are differentially recruited by dissociable decision-making processes – separated in our task across space and time – seem to promote distinct facets of reward-related information and may be uniquely altered between susceptible versus resilient individuals.^77^

Building from this study, future work could explore different stress-related biomarkers, including circuit-specific physiological signatures or neurohormone changes (e.g., cortisol), in response to economic stress.^35,78^ In addition, other two-hit stress models, including early life stress (e.g., maternal separation), followed by economic stress in adulthood could shed light on how different combinations of unique stressors may give rise to distinct stress-related vulnerabilities or pathophysiological states.^79-83^ A limitation of the present study includes controlling economic stress on the task itself. This could be mitigated by transitioning into a reward-scarce environment less abruptly and instead more gradually, as this may lessen the impact on both one’s metabolic demand as well as perceived scarcity. Adjusting resource constraints while in a reward-scarce environment by providing compensated budgets in future studies may also mitigate the effects of economic demand, as others have shown.^84^ Other questions that remain include understanding how economic stress alone could directly promote depression-related phenotypes as is seen in humans and how sex differences may emerge in response to this novel stress model.^85,86^

In summary, we found that using a novel two-hit stress model by combining social stress followed by an economic stressor, mice displayed unique changes in how they recovered from a severe setback in a longitudinal decision-making paradigm. We discovered that mice resilient to social stress were overall more capable in their ability to “bounce back” from an economic setback and did so in a manner that not only ensured food security but also prioritized subjective value. This neuroeconomic task has been translated and validated for use across species including humans, enabling the study of evolutionarily conserved circuits that not only serve basic survival needs but also promote subjective well-being – distinct computational processes that we found are uniquely altered by socioeconomic stress and may be differentially dysfunctional in individuals struggling with stress-related disorders._24,71,87-89_

## Acknowledgments

We thank members of the Nestler and Russo labs for helpful discussion and technical assistance. We thank Caleb Browne and Cindy Aaronson for constructive conversation. We also thank Alexxai Kravitz for assistance in developing the open-source pellet dispensers used in this experiment (www.hackaday.io/project/171116-fed0). Open-source illustrations obtained from SciDraw (www.scidraw.io), credit Federico Claudi.

## Author contributions

Conceptualization: RDC, BMS

Methodology: RDC, SJR, EJN, BMS

Investigation: RDC, FMR, LL, AMT, BMS

Data curation: RDC, BMS

Formal analysis: RDC, ZD, BMS

Visualization: RDC, BMS

Funding acquisition: RDC, DJC, SJR, EJN, BMS

Supervision: RDC, DJC, SJR, EJN, BMS

Writing – original draft: RDC, BMS

Writing – review & editing: all authors

## Data and materials availability

All data is included in the manuscript and/or supporting information.

---

**Supplementary Figure 1.**
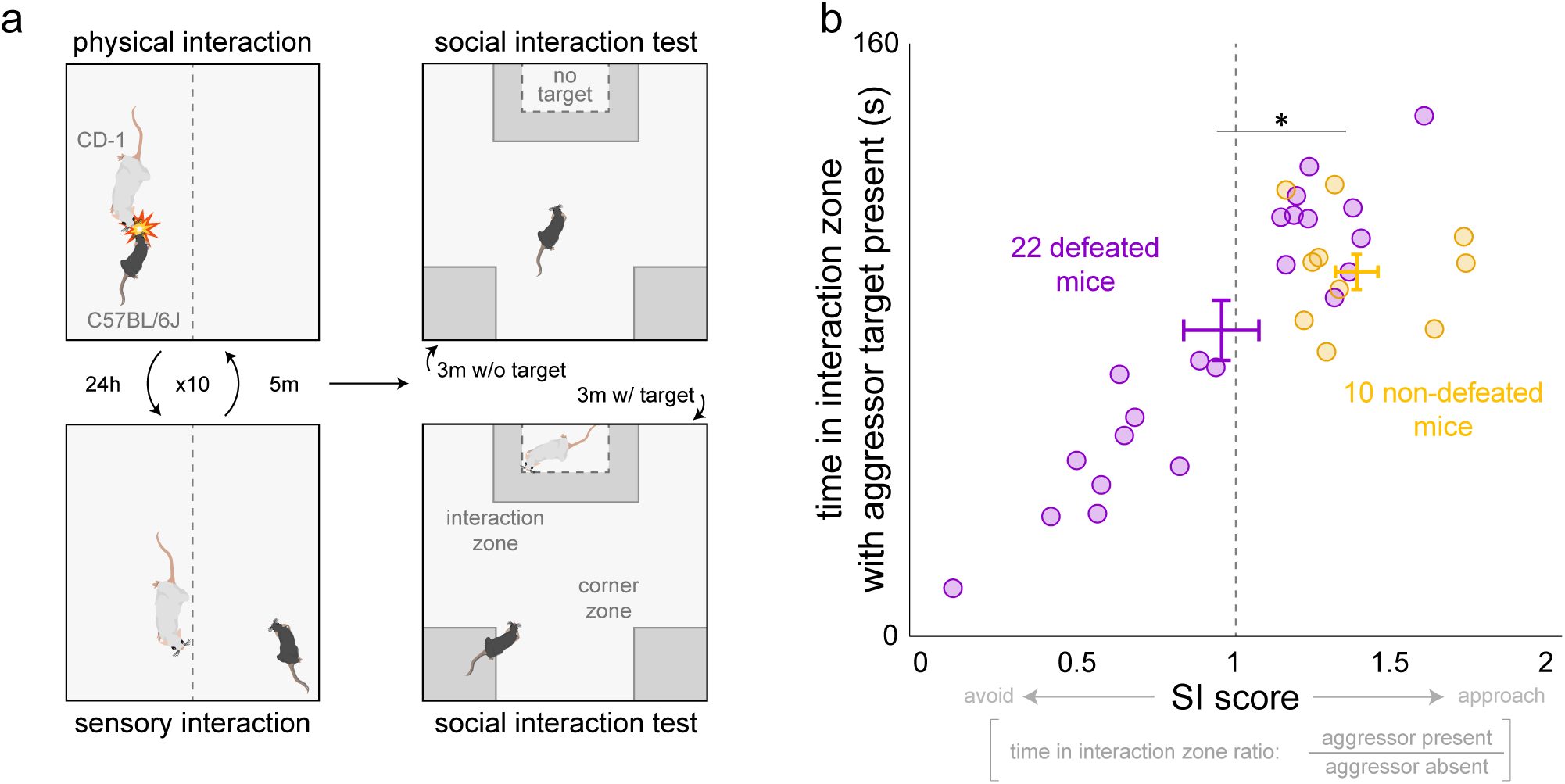
Chronic social defeat stress. (a) Chronic social defeat stress protocol schematic. Following chronic social defeat stress, mice were assayed on the social interaction test. (b) Social avoidance induced by defeat is reflected by a decrease in time spent in the interaction zone when a target CD-1 mouse is present. Vertical dashed gray line indicates an SI score of 1. Dots represent individual animals, x-y error bars ± 1 SEM.

**Supplementary Figure 2.**
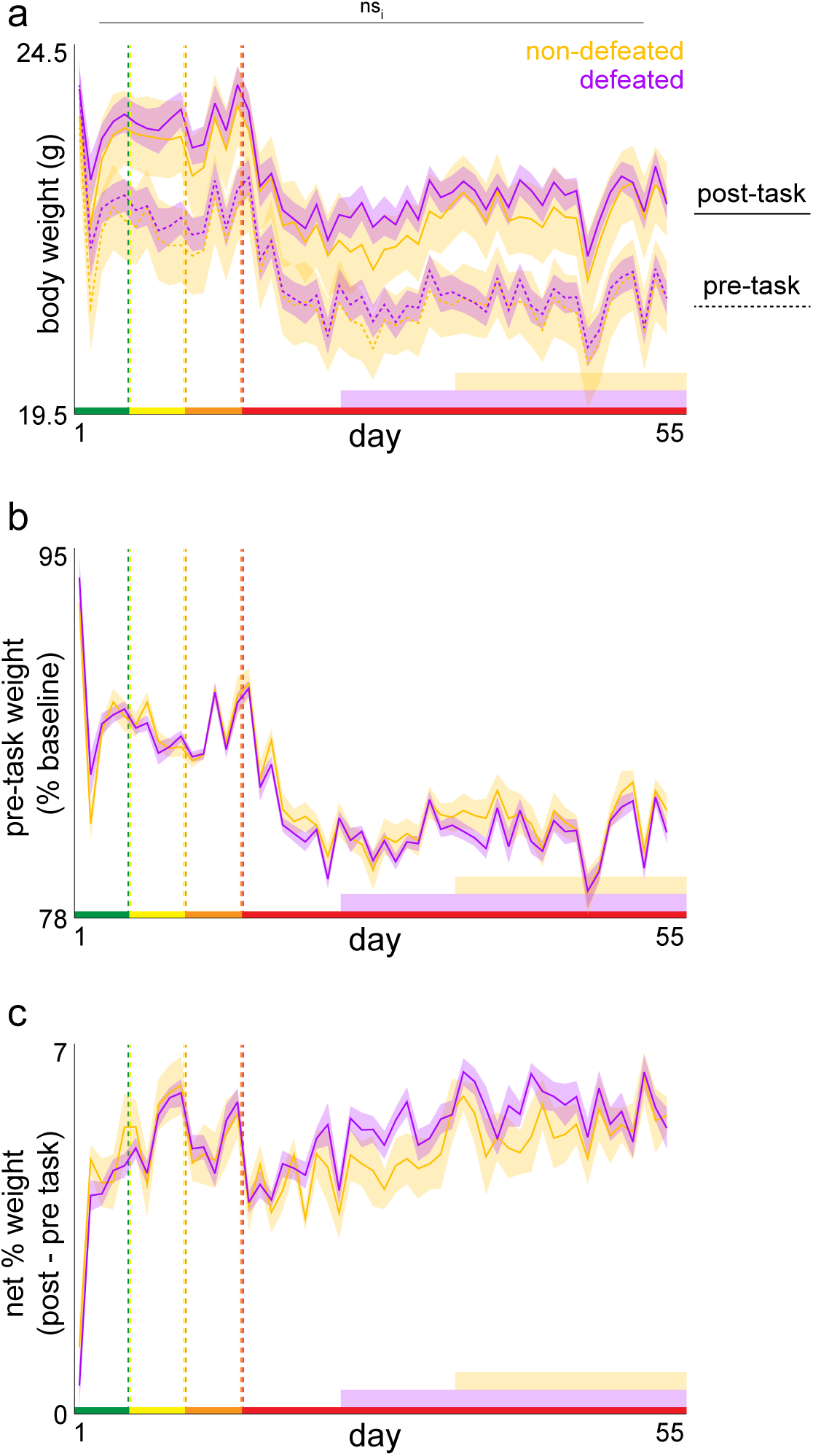
Body weight measures across testing. (a) Average daily body weight measured daily immediately before and after each testing session. (b) Average pre-task weight measured as a % of baseline weights taken 3 days before starting the Restaurant Row task as animals completed the defeat vs. non-defeat protocol while still on ad lib regular chow food before switching to task-earned-only flavored full nutrition pellets. (c) Change in % baseline body weight post minus pre task. No significant interaction between defeated and non-defeated mice across days of testing (pre-task weight: *F*_1,54_=0.965, *p*=0.326; post-task weight: *F*_1,54_=0.051, *p*=0.821). Shaded error bars ± 1 SEM. Not significant interaction (ns_i_).

**Supplementary Figure 3.**
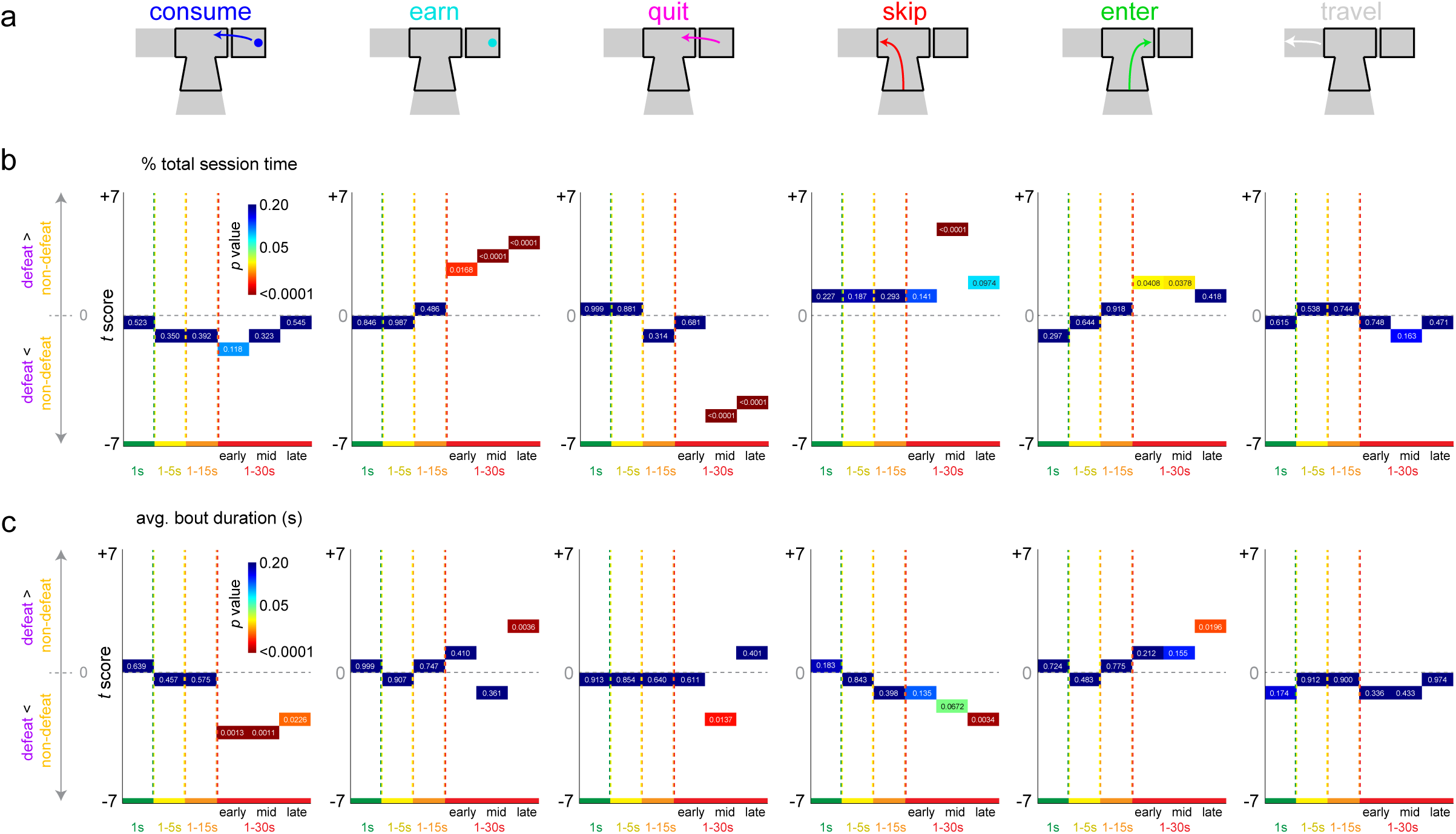
Restructuring of time allocaon across testing. (a) Schematics of all possible and mutually exclusive behaviors mice could be engaged in on the Restaurant Row task. Epochs of testing collapsed into six 5-day bins: 1 s only offers (green epoch, days 1-5), 1-5 s offers (yellow epoch, days 6-10), 1-15s offers (orange epoch, days 11-15), 1-30 s offers (red epoch, three sub-epochs collapsed; early days 16-20; mid days 33-37, late days 51-55). Statistics from unpaired *t* tests plttoed on the y-axis comparing defeat – non-defeat for (b) proportion total session time and (c) average bout duration time metrics. Horizontal dashed gray line indicates *t* score of 0. Heatmap as well as redundant text labels on each data point reflect *p* value.

**Supplementary Figure 4.**
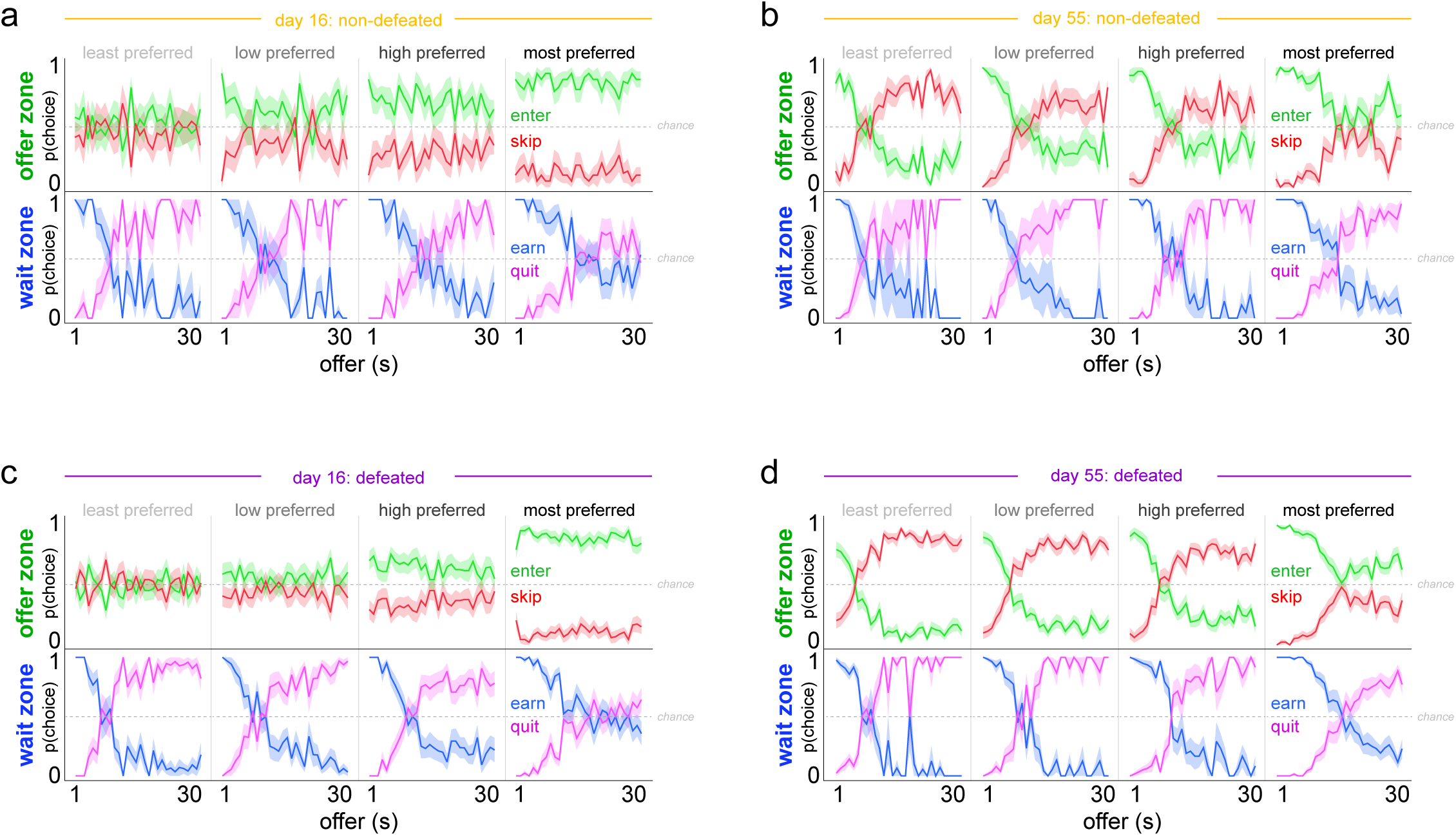
Probability of offer zone and wait zone choices as a function of cost and flavor. The probability of making an enter vs skip choice in the offer zone or an earn vs quit choice in the wait zone was calculated as a function of cued offer length. p(choice) calculated separately in each restaurant ranked from least to most preferred flavors based on end of session number of pellets earned. Data separated by (a-b) non-defeated mice and (c-d) defeated mice taken from the first day in the 1 to 30 s reward-scarce environment (day 16, a, c) and the last day in this experiment (day 55, b, d). Note that on day 16, in the offer zone, mice made enter vs skip decisions without factoring in cued offer cost but sll displayed biases to enter depending on the ordinal ranking of each restaurant. In contrast, by day 55, experienced mice made enter vs skip decisions in the offer zone not only based on the ordinal ranking of each restaurant but also as a function of cued offer cost. The fact that experienced mice treat the same tones differently in each restaurant indicates mice integrate auditory information with spaal information. Horizontal dashed gray line indicates chance p(choice) at 0.5. Shaded error bars ± 1 SEM.

**Supplementary Figure 5.**
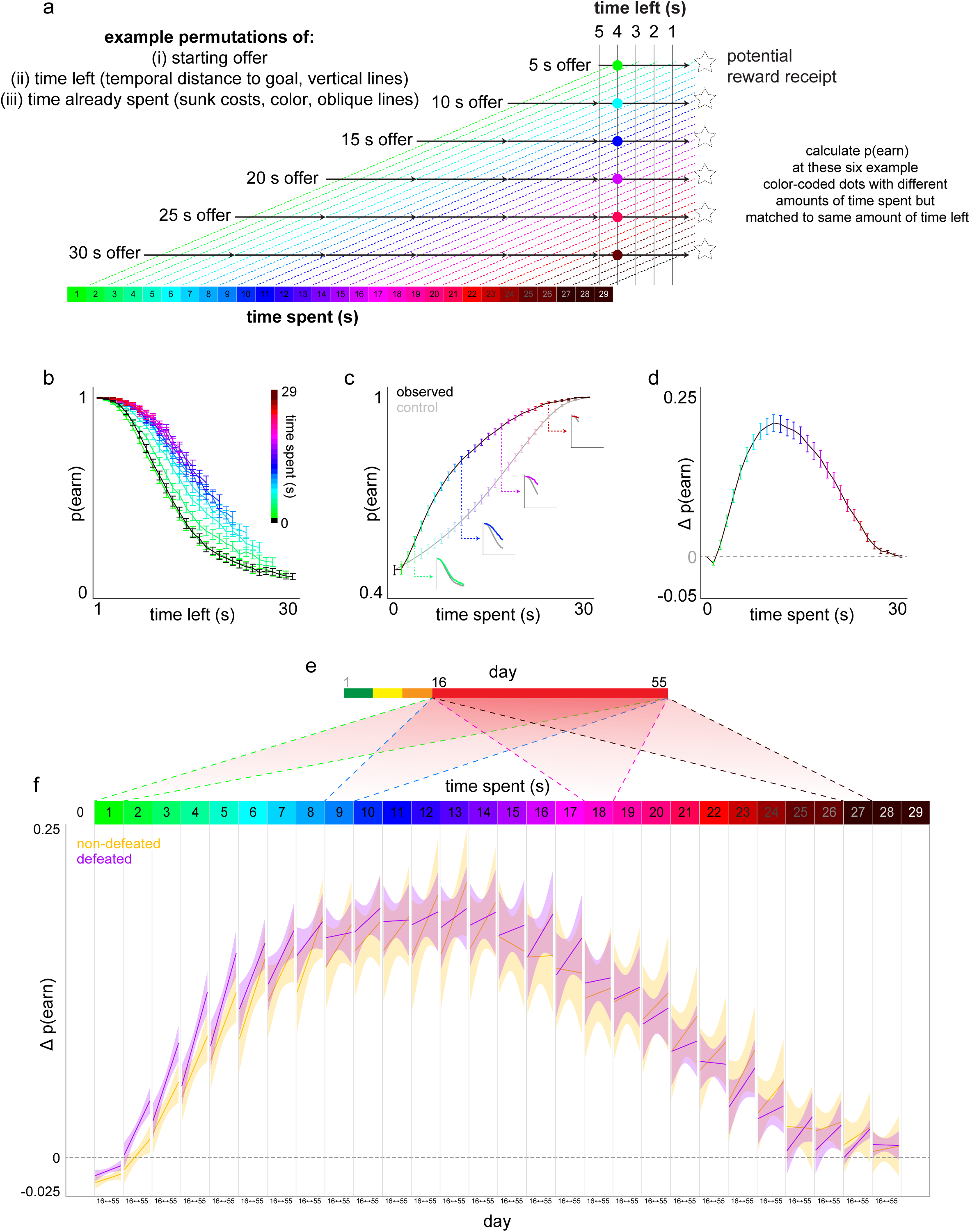
Visual explanation of the sunk cost analysis with sensitivity rates across testing. (a) All possible trial combinations of [me spent, time left] pairs can be compiled based on the randomly presented starting off costs and by using analysis points at progressively longer wait mes during the countdown in the wait zone. Displayed here are 6 example starng offers (5, 10, 15, 20, 25, and 30 s offers, although this analysis extends to all 1 to 30 s offers). Colored diagonal dashed lines represent increasing amounts of time already spent waiting in the wait zone as animals progress toward earning a reward (dashed star shapes). 5 vertical solid blank lines reflect an example epoch of the last 5 s remaining in the countdown before potentially obtaining a reward common to all starting offer lengths. Using the 4 s remaining vertical solid black line as an example, the 6 color-coded dots indicate separate sunk cost conditions with increasing amounts of time already spent (colors) matched for the same temporal distance to the goal (4 s in this example). Of the trials in which the mouse had not yet quit by this point (i.e., trials “still live”), the probably of staying in the wait zone and earning a reward can be calculated at this analysis point. Sensitivity to sunk costs is represented by an increase in p(earn) as a function of time already waited, or an escalation of commitment. This effect is observed independent of and orthogonal to temporal distance to the goal because this analysis can be repeated at every instance of “time left” and a conserved effect of sunk costs on increasing p(earn) can be observed. By collapsing across the time left dimension and only focusing on the time spent (color) dimension, we can appreciate a change in p(earn) relave to the same amount of titime left for all 0 s sunk cost condions. (b-d) Data from the 1 to 30 s epoch across all 32 mice used to demonstrate analysis steps. (b) p(earn) in the wait zone calculated as a function of time left in the countdown (x-axis) and time spent (color). Note p(earn) increases as a function of time spent regardless of temporal distance to the goal. (c) p(earn) data from (b) collapsed across time left in order to capture envelope of sensivitity to sunk costs. Control analysis iteravely collapses the 0 s sunk condition (black curve) repeatedly matched to the same number of time left datapoints for a given sunk cost condition. Small inset graphs depict four examples from (b) collapsed into single datapoints in (c) with matching number of truncated datapoints in the 0 s sunk condition collapsed into the gray control curve. (d) The magnitude of sensivitity to sunk costs, Δp(earn), is calculated by subtracting the observed – control curves in (c). (e) This analysis was performed across the entire testing epoch (see main figure 4g-h) however here we focus just on the 1 to 30 s reward scarce environment (days 16 to 55) in order to display the change in p(earn) for each sunk cost condition across days of testing in (f). The x-axis in (f) reflects days 16 to 55 repeated for each sunk cost condition (four example shadow boxes from (e) show the reward-scarce epoch [red] compressed into each sunk cost condition). Linear fits across the entire day 16 to 55 epoch were performed for each sunk cost condition. Note that the rate at which mice acquire sensivitity to sunk costs across days of testing is enhanced in defeated mice particularly during the initial few seconds during re-evaluation change-of-mind decisions. Error bars ± 1 SEM. Shaded error bars 95% CI of linear fits.

**Supplementary Figure 6.**
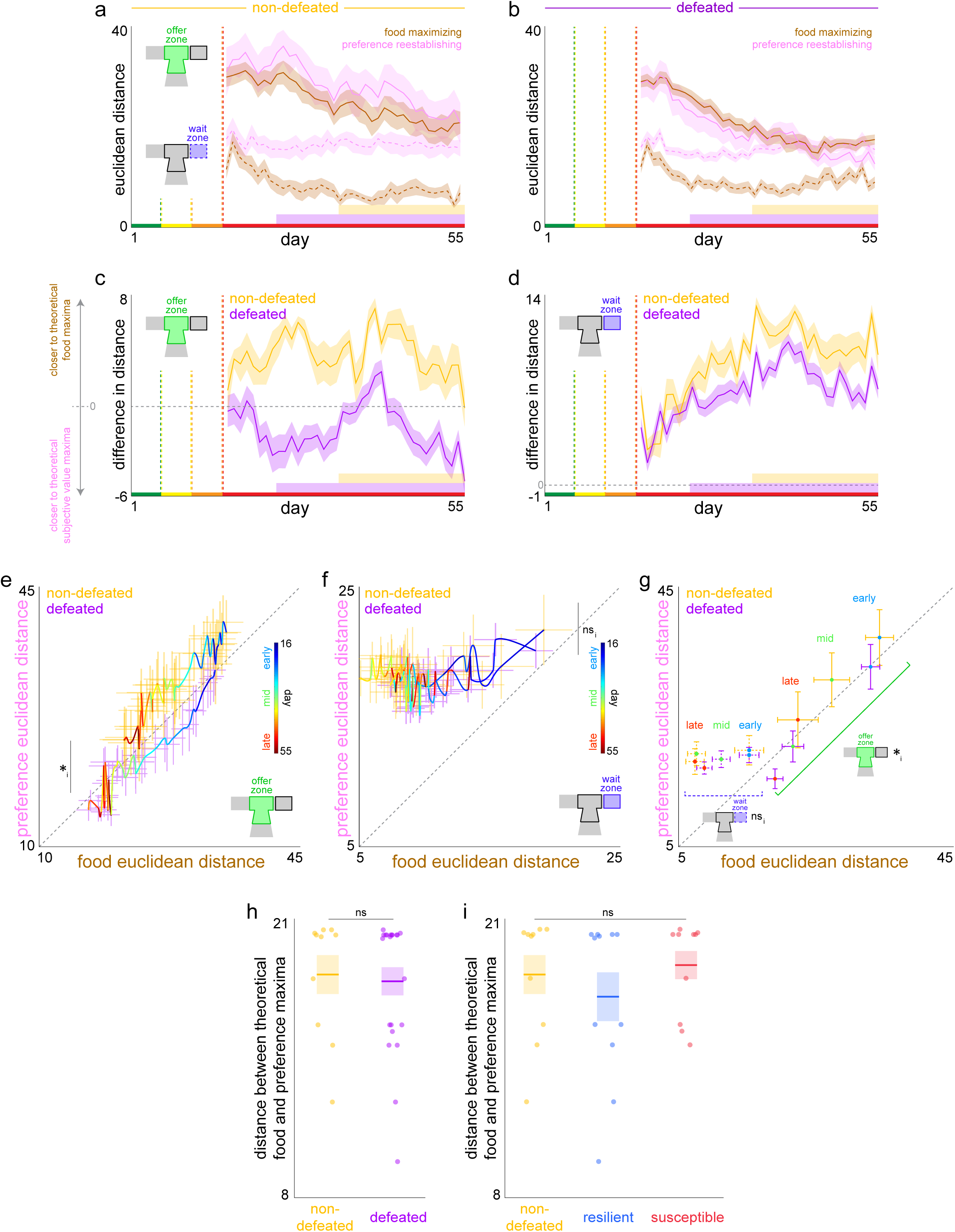
Decision policy Euclidean distances. (a-b) A redisplay of Euclidean distances measured across days of testing for both food maximizing and preference reestablishing decision policies in both the offer zone (solid) and wait zone (dashed) for (a) non-defeated mice and (b) defeated mice. Note: in the wait zone, preference reestablishing Euclidean distances (pink dashed lines) do not significantly change across days of testing for either group of mice (*F*_1,39_=2.415, *p*=0.121). (c-d) Euclidean distance difference scores in the (c) offer zone and (d) wait zone calculated by subtracting food maximizing distances from preference reestablishing distances (thus, a more negative difference score equates to a decision policy that is closer to the preference reestablishing than food theoretical maxima). Horizontal dashed gray line indicates a difference score of 0. Note: in the OZ, defeated mice overall have a difference score below zero (sign test: *t*=-5.099, *p*<0.0001) while non-defeated mice overall have a difference score above zero (sign test: *t*=+9.647, *p*<0.0001). (e-g) Redisplay of food by preference Euclidean distance decision policy trajectories between defeated and non-defeated mice across days of testing (color heat map line) in the (e) offer zone and (f) wait zone. X-Y error bars reflect each single-day standard error of the mean. (g) Summary of data from (e-f) for both the offer zone (solid) and wait zone (dashed) that also appear in main figure 5g-h where averages are taken from early (days 16-20, blue), mid (days 34-38, green), and late (days 51-55, red) of testing. Note: groups of mice did not differ in either distance in either zone during the early time period (OZ – food: *F*_1,4_=0.386, *p*=0.539; preference: *F*_1,4_=0.782, *p*=0.384; WZ – food: *F*_1,4_=0.001, *p*=0.996; preference: *F*_1,4_=0.156, *p*=0.696). (h-i) No significant differences in distances between theoretical food maximum and theoretical subjective “preference” value maximum among groups of mice (h, defeat vs non-defeated groups, *F*_1,31_=0.064, *p*=0.802; i, including split by resilient and susceptible subgroups, *F*_2,31_=0.697, *p*=0.506). Shaded and x-y error bars ± 1 SEM. Not significant interaction (ns_i_).

## References

1. American Psychiatric Association., American Psychiatric Association. DSM-5 Task Force. Diagnostic and statistical manual of mental disorders : DSM-5. 5th ed. Washington, D.C.: American Psychiatric Association; 2013.

2. Feder A, Nestler EJ, Charney DS. Psychobiology and molecular genetics of resilience. Nat Rev Neurosci 2009; 10(6): 446–57.

3. Charney DS. Psychobiological mechanisms of resilience and vulnerability: implications for successful adaptation to extreme stress. Am J Psychiatry 2004; 161(2): 195–216.

4. Feder A, Fred-Torres S, Southwick SM, Charney DS. The Biology of Human Resilience: Opportunities for Enhancing Resilience Across the Life Span. Biol Psychiatry 2019; 86(6): 443–53.

5. Han MH, Nestler EJ. Neural Substrates of Depression and Resilience. Neurotherapeutics 2017; 14(3): 677–86.

6. Russo SJ, Murrough JW, Han MH, Charney DS, Nestler EJ. Neurobiology of resilience. Nat Neurosci 2012; 15(11): 1475–84.

7. Loewenstein G, Rick S, Cohen JD. Neuroeconomics. Annu Rev Psychol 2008; 59: 647–72.

8. Camerer CF. Neuroeconomics: opening the gray box. Neuron 2008; 60(3): 416–9.

9. Montague PR. Neuroeconomics: a view from neuroscience. Funct Neurol 2007; 22(4): 219–34.

10. Glimcher PW, Rustichini A. Neuroeconomics: the consilience of brain and decision. Science 2004; 306(5695): 447–52.

11. Price RH, Choi JN, Vinokur AD. Links in the chain of adversity following job loss: how financial strain and loss of personal control lead to depression, impaired functioning, and poor health. J Occup Health Psychol 2002; 7(4): 302–12.

12. Robinson OJ, Bond RL, Roiser JP. The impact of stress on financial decision-making varies as a function of depression and anxiety symptoms. PeerJ 2015; 3: e770.

13. Kalenscher T, van Wingerden M. Why we should use animals to study economic decision making - a perspective. Front Neurosci 2011; 5: 82.

14. Liem R, Liem J. Social class and mental illness reconsidered: the role of economic stress and social support. J Health Soc Behav 1978; 19(2): 139–56.

15. MacFadyen A, MacFadyen H, Prince N. Economic stress and psychological well-being: An economic psychology framework. J Economic Psychology 1996; 17(3): 291–311.

16. Perzow SED, Bray BC, Wadsworth ME. Financial stress response profiles and psychosocial functioning in low-income parents. J Fam Psychol 2018; 32(4): 517–27.

17. Krishnan V, Han MH, Graham DL, et al. Molecular adaptations underlying susceptibility and resistance to social defeat in brain reward regions. Cell 2007; 131(2): 391–404.

18. Sweis BM, Thomas MJ, Redish AD. Mice learn to avoid regret. PLoS Biol 2018; 16(6): e2005853.

19. Steiner AP, Redish AD. Behavioral and neurophysiological correlates of regret in rat decision-making on a neuroeconomic task. Nat Neurosci 2014; 17(7): 995–1002.

20. Durand-de Cuttoli R, Martinez-Rivera FJ, Li L, et al. Distinct forms of regret linked to resilience versus susceptibility to stress are regulated by region-specific CREB function in mice. Science Advances 2022; 8(42): eadd5579.

21. Sweis BM, Larson EB, Redish AD, Thomas MJ. Altering gain of the infralimbic-to-accumbens shell circuit alters economically dissociable decision-making algorithms. Proc Natl Acad Sci U S A 2018; 115(27): E6347–E55.

22. Sweis BM, Redish AD, Thomas MJ. Prolonged abstinence from cocaine or morphine disrupts separable valuations during decision conflict. Nat Commun 2018; 9(1): 2521.

23. Arkes H, Blumer C. The psychology of sunk cost. Organizational Behavior and Human Decision Processes 1985; 35(1): 124–40.

24. Sweis BM, Abram SV, Schmidt BJ, et al. Sensitivity to “sunk costs” in mice, rats, and humans. Science 2018; 361(6398): 178–81.

25. Redish AD, Abram SV, Cunningham PJ, et al. Sunk cost sensitivity during change-of-mind decisions is informed by both the spent and remaining costs. Commun Biol 2022; 5(1): 1337.

26. Falagas ME, Vouloumanou EK, Mavros MN, Karageorgopoulos DE. Economic crises and mortality: a review of the literature. Int J Clin Pract 2009; 63(8): 1128–35.

27. Ss C, d G, Ja S, Th L, At C. Was the economic crisis 1997-1998 responsible for rising suicide rates in East/Southeast Asia? A time-trend analysis for Japan, Hong Kong, South Korea, Taiwan, Singapore and Thailand. Soc Sci Med 2009; 68(7): 1322–31.

28. Houdmont J, Kerr R, Addley K. Psychosocial factors and economic recession: the Stormont Study. Occup Med (Lond) 2012; 62(2): 98–104.

29. Takahashi T. Neuroeconomics of suicide. Neuro Endocrinol Lett 2011; 32(4): 400–4.

30. Jenkins R, Bhugra D, Bebbington P, et al. Debt, income and mental disorder in the general population. Psychol Med 2008; 38(10): 1485–93.

31. K P, K M. Unemployment impairs mental health: Meta-analyses. J Vocat Behav 2009; 74(3): 264–82.

32. Giorgi G, Arcangeli G, Mucci N, Cupelli V. Economic stress in the workplace: The impact of fear of the crisis on mental health. Work 2015; 51(1): 135–42.

33. Gonda X, Petschner P, Eszlari N, et al. Effects of Different Stressors Are Modulated by Different Neurobiological Systems: The Role of GABA-A Versus CB1 Receptor Gene Variants in Anxiety and Depression. Front Cell Neurosci 2019; 13: 138.

34. Gonda X, Eszlari N, Kovacs D, et al. Financial difficulties but not other types of recent negative life events show strong interactions with 5-HTTLPR genotype in the development of depressive symptoms. Transl Psychiatry 2016; 6(5): e798.

35. Sripada RK, Swain JE, Evans GW, Welsh RC, Liberzon I. Childhood poverty and stress reactivity are associated with aberrant functional connectivity in default mode network. Neuropsychopharmacology 2014; 39(9): 2244–51.

36. Trivedi MH, Rush AJ, Wisniewski SR, et al. Evaluation of outcomes with citalopram for depression using measurement-based care in STAR*D: implications for clinical practice. Am J Psychiatry 2006; 163(1): 28–40.

37. Spellman T, Svei M, Kaminsky J, Manzano-Nieves G, Liston C. Prefrontal deep projection neurons enable cognitive flexibility via persistent feedback monitoring. Cell 2021; 184(10): 2750–66 e17.

38. Hasz BM, Redish AD. Dorsomedial prefrontal cortex and hippocampus represent strategic context even while simultaneously changing representation throughout a task session. Neurobiol Learn Mem 2020; 171: 107215.

39. Massar SA, Lim J, Sasmita K, Chee MW. Rewards boost sustained attention through higher effort: A value-based decision making approach. Biol Psychol 2016; 120: 21–7.

40. Winstanley CA, Floresco SB. Deciphering Decision Making: Variation in Animal Models of Effort- and Uncertainty-Based Choice Reveals Distinct Neural Circuitries Underlying Core Cognitive Processes. J Neurosci 2016; 36(48): 12069–79.

41. Studer B, Koch C, Knecht S, Kalenscher T. Conquering the inner couch potato: precommitment is an effective strategy to enhance motivation for effortful actions. Philos Trans R Soc Lond B Biol Sci 2019; 374(1766): 20180131.

42. Kalenscher T, Pennartz CM. Is a bird in the hand worth two in the future? The neuroeconomics of intertemporal decision-making. Prog Neurobiol 2008; 84(3): 284–315.

43. Der-Avakian A, Mazei-Robison MS, Kesby JP, Nestler EJ, Markou A. Enduring deficits in brain reward function after chronic social defeat in rats: susceptibility, resilience, and antidepressant response. Biol Psychiatry 2014; 76(7): 542–9.

44. Yoshida K, Drew MR, Kono A, Mimura M, Takata N, Tanaka KF. Chronic social defeat stress impairs goal-directed behavior through dysregulation of ventral hippocampal activity in male mice. Neuropsychopharmacology 2021; 46(9): 1606–16.

45. Riga D, Theijs JT, De Vries TJ, Smit AB, Spijker S. Social defeat-induced anhedonia: effects on operant sucrose-seeking behavior. Front Behav Neurosci 2015; 9: 195.

46. Sweis BM, Veverka KK, Dhillon ES, Urban JH, Lucas LR. Individual differences in the effects of chronic stress on memory: behavioral and neurochemical correlates of resiliency. Neuroscience 2013; 246: 142–59.

47. Ordones Sanchez E, Bavley CC, Deutschmann AU, et al. Early life adversity promotes resilience to opioid addiction-related phenotypes in male rats and sex-specific transcriptional changes. Proc Natl Acad Sci U S A 2021; 118(8).

48. Bhatnagar S. Rethinking stress resilience. Trends Neurosci 2021; 44(12): 936–45.

49. Haynes S, Lacagnina A, Seong HS, et al. CRF neurons establish resilience via stress-history dependent BNST modulation. biorxiv 2022: 2022.08.31.505596.

50. Larrieu T, Cherix A, Duque A, et al. Hierarchical Status Predicts Behavioral Vulnerability and Nucleus Accumbens Metabolic Profile Following Chronic Social Defeat Stress. Curr Biol 2017; 27(14): 2202–10 e4.

51. Patterson ZR, Khazall R, Mackay H, Anisman H, Abizaid A. Central ghrelin signaling mediates the metabolic response of C57BL/6 male mice to chronic social defeat stress. Endocrinology 2013; 154(3): 1080–91.

52. Chuang JC, Cui H, Mason BL, et al. Chronic social defeat stress disrupts regulation of lipid synthesis. J Lipid Res 2010; 51(6): 1344–53.

53. Razzoli M, Frontini A, Gurney A, et al. Stress-induced activation of brown adipose tissue prevents obesity in conditions of low adaptive thermogenesis. Mol Metab 2016; 5(1): 19–33.

54. Lutter M, Elmquist J. Depression and metabolism: linking changes in leptin and ghrelin to mood. F1000 Biol Rep 2009; 1: 63.

55. Duin AA, Aman L, Schmidt B, Redish AD. Certainty and uncertainty of the future changes planning and sunk costs. Behav Neurosci 2021.

56. Pompilio L, Kacelnik A, Behmer ST. State-dependent learned valuation drives choice in an invertebrate. Science 2006; 311(5767): 1613–5.

57. Zentall TR. Within-trial contrast: when you see it and when you don’t. Learn Behav 2008; 36(1): 19-22; discussion 3-8.

58. Wikenheiser AM, Stephens DW, Redish AD. Subjective costs drive overly patient foraging strategies in rats on an intertemporal foraging task. Proc Natl Acad Sci U S A 2013; 110(20): 8308–13.

59. Murra D, Hilde KL, Fitzpatrick A, Maras PM, Watson SJ, Akil H. Characterizing the Behavioral and Neuroendocrine Features of Susceptibility and Resilience to Social Stress. bioRxiv 2021: 2021.12.13.472392.

60. Lorsch ZS, Hamilton PJ, Ramakrishnan A, et al. Stress resilience is promoted by a Zfp189-driven transcriptional network in prefrontal cortex. Nat Neurosci 2019; 22(9): 1413–23.

61. Willmore L, Cameron C, Yang J, Witten IB, Falkner AL. Behavioural and dopaminergic signatures of resilience. Nature 2022; 611(7934): 124–32.

62. Selye H. The stress concept. Can Med Assoc J 1976; 115(8): 718.

63. Koolhaas JM, Bartolomucci A, Buwalda B, et al. Stress revisited: a critical evaluation of the stress concept. Neurosci Biobehav Rev 2011; 35(5): 1291–301.

64. Nasca C, Menard C, Hodes G, et al. Multidimensional Predictors of Susceptibility and Resilience to Social Defeat Stress. Biol Psychiatry 2019; 86(6): 483–91.

65. Ball TM, Gunaydin LA. Measuring maladaptive avoidance: from animal models to clinical anxiety. Neuropsychopharmacology 2022; 47(5): 978–86.

66. Calipari ES, Siciliano CA, Zimmer BA, Jones SR. Brief intermittent cocaine self-administration and abstinence sensitizes cocaine effects on the dopamine transporter and increases drug seeking. Neuropsychopharmacology 2015; 40(3): 728–35.

67. Farris SG, Aston ER, Abrantes AM, Zvolensky MJ. Tobacco demand, delay discounting, and smoking topography among smokers with and without psychopathology. Drug Alcohol Depend 2017; 179: 247–53.

68. Farris SG, Aston ER, Zvolensky MJ, Abrantes AM, Metrik J. Psychopathology and tobacco demand. Drug Alcohol Depend 2017; 177: 59–66.

69. Murphy JG, Yurasek AM, Dennhardt AA, et al. Symptoms of depression and PTSD are associated with elevated alcohol demand. Drug Alcohol Depend 2013; 127(1-3): 129–36.

70. Salamone, Correa, Yohn, et al. Behavioral activation, effort-based choice, and elasticity of demand for motivational stimuli: Basic and translational neuroscience approaches. Motivation Science 2017; 3(3): 208–29.

71. Abram SV, Hanke M, Redish AD, MacDonald AW, 3rd. Neural signatures underlying deliberation in human foraging decisions. Cogn Affect Behav Neurosci 2019; 19(6): 1492–508.

72. Diehl G, Redish AD. Differential processing of decision information in subregions of rodent medial prefrontal cortex. biorxiv 2022.

73. Johnson A, Redish AD. Neural ensembles in CA3 transiently encode paths forward of the animal at a decision point. J Neurosci 2007; 27(45): 12176–89.

74. Schmidt B, Duin AA, Redish AD. Disrupting the medial prefrontal cortex alters hippocampal sequences during deliberative decision making. J Neurophysiol 2019; 121(6): 1981–2000.

75. Schmidt B, Redish AD. Disrupting the medial prefrontal cortex with designer receptors exclusively activated by designer drug alters hippocampal sharp-wave ripples and their associated cognitive processes. Hippocampus 2021.

76. Stone C, Mattingley JB, Rangelov D. On second thoughts: changes of mind in decision-making. Trends Cogn Sci 2022; 26(5): 419–31.

77. Sweis BM, Thomas MJ, Redish AD. Beyond simple tests of value: measuring addiction as a heterogeneous disease of computation-specific valuation processes. Learn Mem 2018; 25(9): 501–12.

78. Dowd JB, Simanek AM, Aiello AE. Socio-economic status, cortisol and allostatic load: a review of the literature. Int J Epidemiol 2009; 38(5): 1297–309.

79. Pena CJ, Kronman HG, Walker DM, et al. Early life stress confers lifelong stress susceptibility in mice via ventral tegmental area OTX2. Science 2017; 356(6343): 1185–8.

80. Davis SM, Rice M, Rudlong J, Eaton V, King T, Burman MA. Neonatal pain and stress disrupts later-life pavlovian fear conditioning and sensory function in rats: Evidence for a two-hit model. Dev Psychobiol 2018; 60(5): 520–33.

81. Faraji J, Soltanpour N, Ambeskovic M, et al. Evidence for Ancestral Programming of Resilience in a Two-Hit Stress Model. Front Behav Neurosci 2017; 11: 89.

82. Jaric I, Rocks D, Cham H, Herchek A, Kundakovic M. Sex and Estrous Cycle Effects on Anxiety- and Depression-Related Phenotypes in a Two-Hit Developmental Stress Model. Front Mol Neurosci 2019; 12: 74.

83. Neumaier JF, Edwards E, Plotsky PM. 5-HT(1B) mrna regulation in two animal models of altered stress reactivity. Biol Psychiatry 2002; 51(11): 902–8.

84. van Wingerden M, Marx C, Kalenscher T. Budget Constraints Affect Male Rats’ Choices between Differently Priced Commodities. PLoS One 2015; 10(6): e0129581.

85. Schneider B, Richard J, Younger A, Freeman P. A longitudinal exploration of the continuity of children’s social participation and social withdrawal across socioeconomic status levels and social settings. European Journal of Social Psychology 2000; 30(4): 497–519.

86. Lewis Brown R, Richman JA. Sex differences in mediating and moderating processes linking economic stressors, psychological distress, and drinking. J Stud Alcohol Drugs 2012; 73(5): 811–9.

87. Abram SV, Breton YA, Schmidt B, Redish AD, MacDonald AW, 3rd. The Web-Surf Task: A translational model of human decision-making. Cogn Affect Behav Neurosci 2016; 16(1): 37–50.

88. Abram SV, Redish AD, MacDonald AW, 3rd. Learning From Loss After Risk: Dissociating Reward Pursuit and Reward Valuation in a Naturalistic Foraging Task. Front Psychiatry 2019; 10: 359.

89. Redish AD, Kepecs A, Anderson LM, et al. Computational validity: using computation to translate behaviours across species. Philos Trans R Soc Lond B Biol Sci 2022; 377(1844): 20200525.

