## Supplementary Figures for "A double-hit of social and economic stress in mice precipitates changes in decision-making strategies"

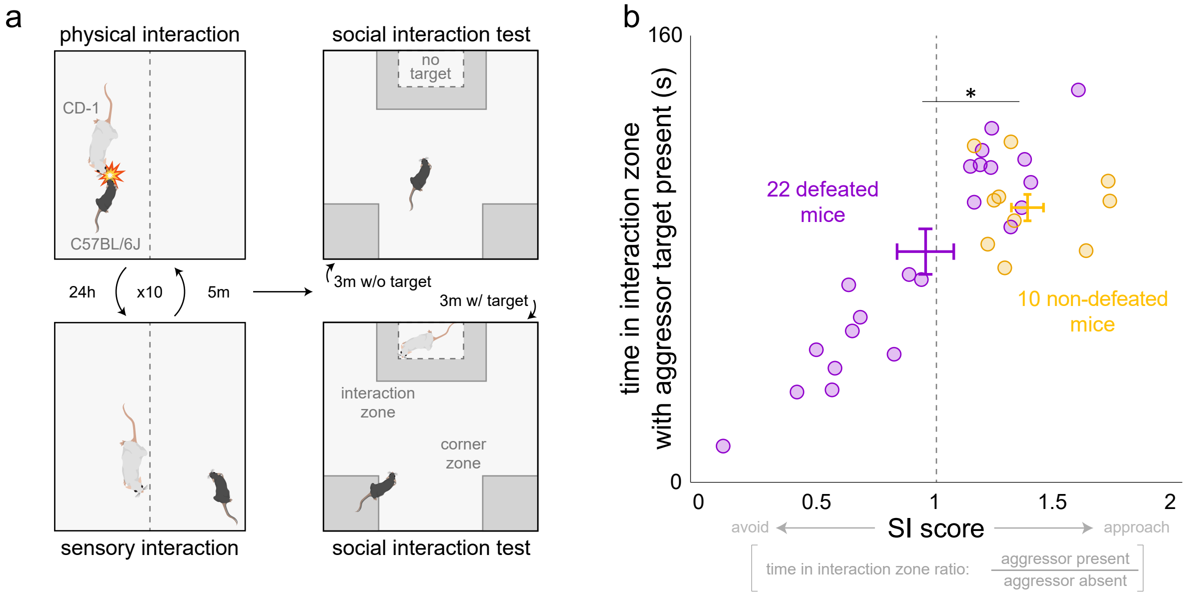


**Supplementary Figure 1. Chronic social defeat stress.** (a) Chronic social defeat stress protocol schematic. Following chronic social defeat stress, mice were assayed on the social interaction test. (b) Social avoidance induced by defeat is reflected by a decrease in time spent in the interaction zone when a target CD-1 mouse is present. Vertical dashed gray line indicates an SI score of 1. Dots represent individual animals, x-y error bars ± 1 SEM.

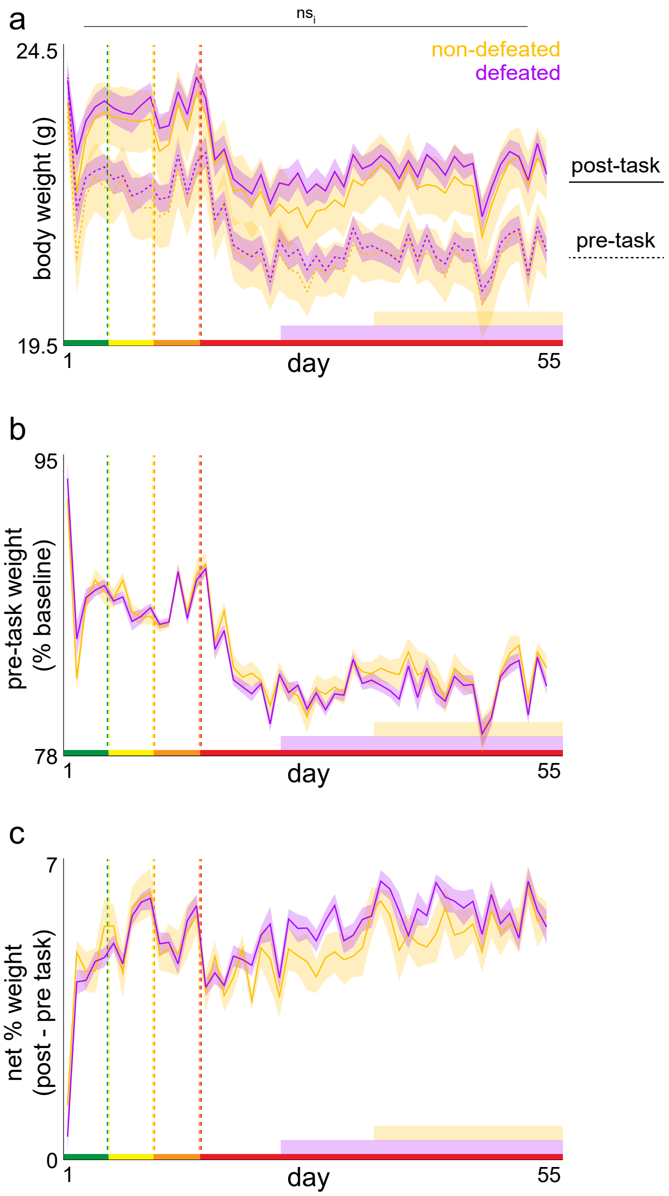


**Supplementary Figure 2. Body weight measures across testing.** (a) Average daily body weight measured daily immediately before and after each testing session. (b) Average pre-task weight measured as a % of baseline weights taken 3 days before starting the Restaurant Row task as animals completed the defeat vs. non-defeat protocol while still on ad lib regular chow food before switching to task-earned-only flavored full nutrition pellets. (c) Change in % baseline body weight post minus pre task. No significant interaction between defeated and non-defeated mice across days of testing (pre-task weight: $F_{1,54}$=0.965, $p$=0.326; post-task weight: $F_{1,54}$=0.051, $p$=0.821). Shaded error bars ± 1 SEM. Not significant interaction (ns$_i$).

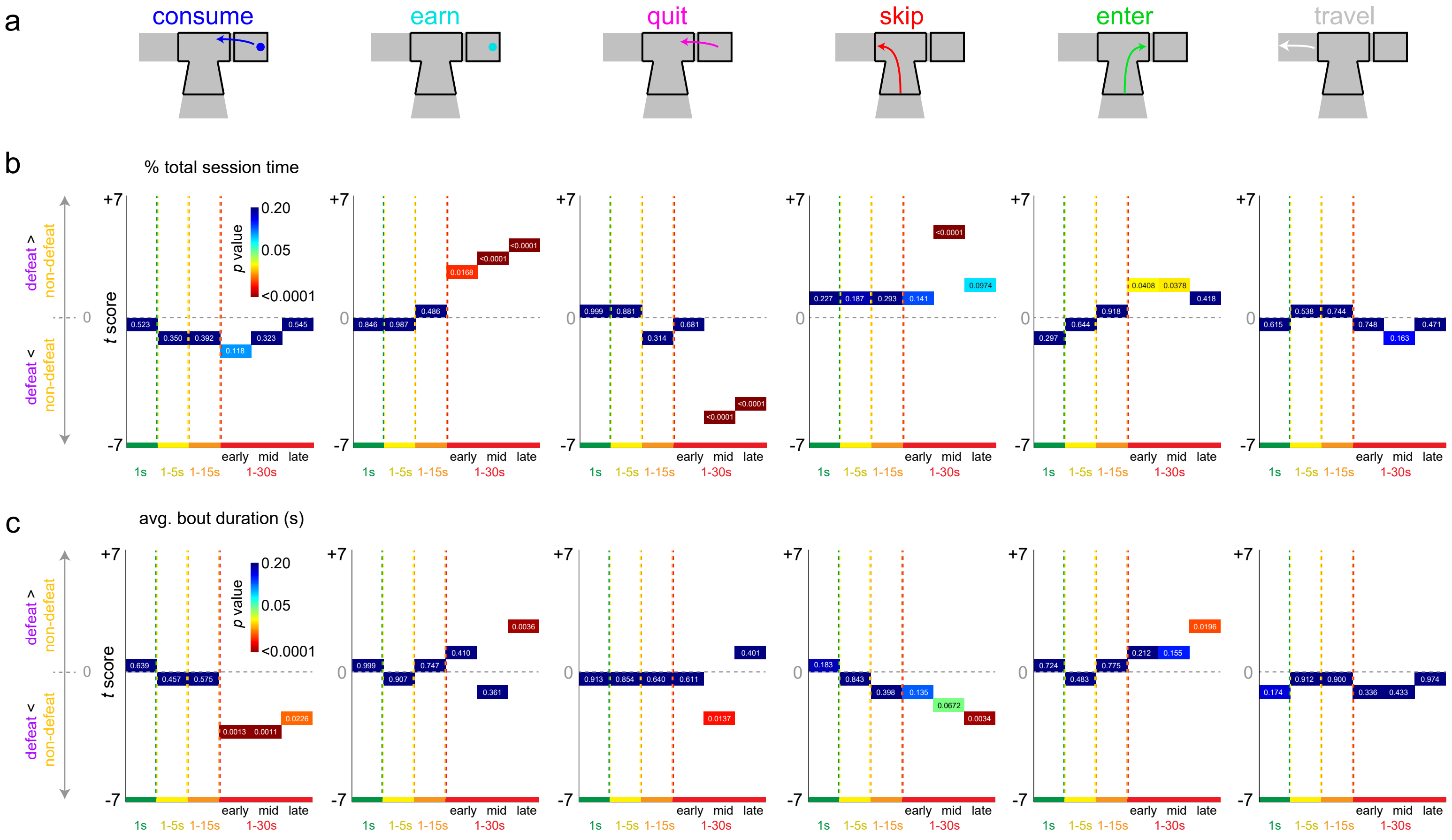


**Supplementary Figure 3. Restructuring of time allocation across testing.** (a) Schematics of all possible and mutually exclusive behaviors mice could be engaged in on the Restaurant Row task. Epochs of testing collapsed into six 5-day bins: 1 s only offers (green epoch, days 1-5), 1-5 s offers (yellow epoch, days 6-10), 1-15s offers (orange epoch, days 11-15), 1-30 s offers (red epoch, three sub-epochs collapsed; early days 16-20; mid days 33-37, late days 51-55). Statistics from unpaired *t* tests plotted on the y-axis comparing defeat – non-defeat for (b) proportion total session time and (c) average bout duration time metrics. Horizontal dashed gray line indicates *t* score of 0. Heatmap as well as redundant text labels on each data point reflect *p* value.

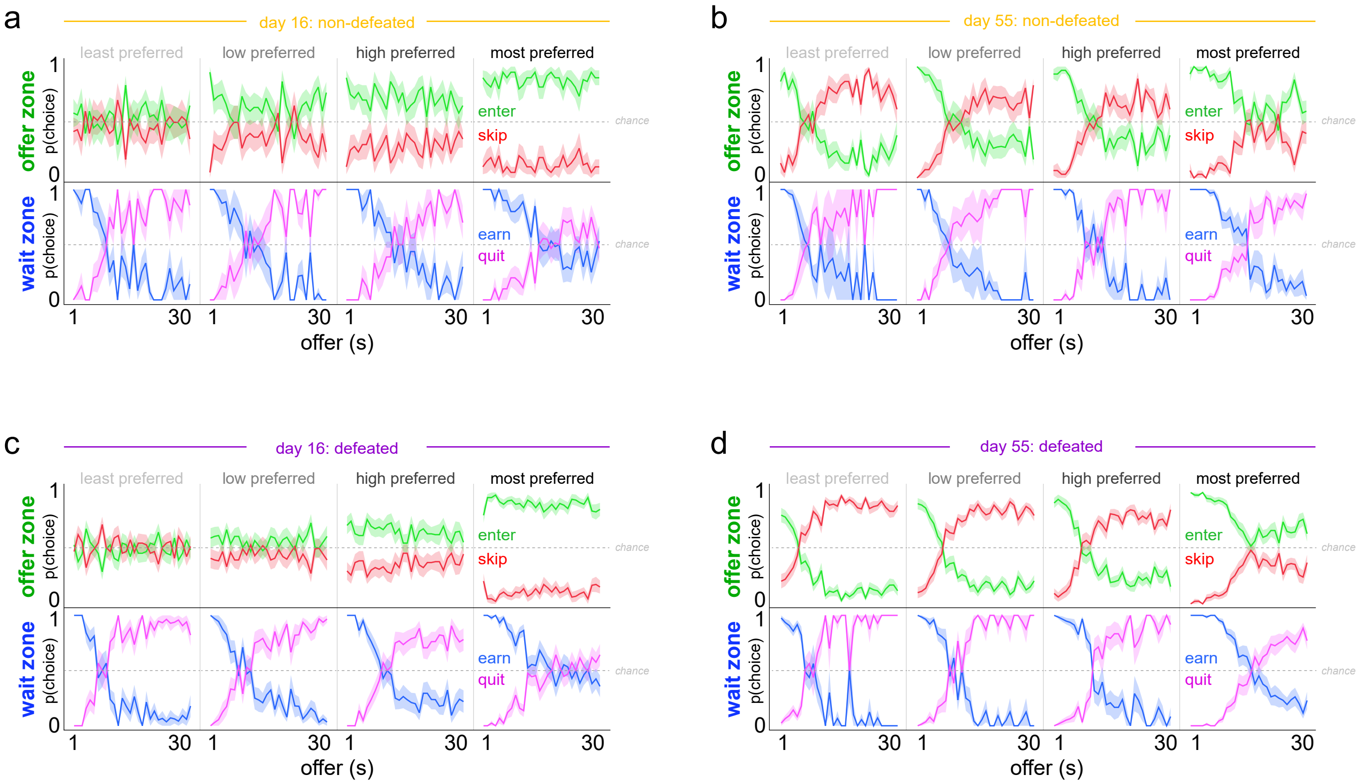


**Supplementary Figure 4. Probability of offer zone and wait zone choices as a function of cost and flavor.** The probability of making an enter vs skip choice in the offer zone or an earn vs quit choice in the wait zone was calculated as a function of cued offer length. p(choice) calculated separately in each restaurant ranked from least to most preferred flavors based on end of session number of pellets earned. Data separated by (a-b) non-defeated mice and (c-d) defeated mice taken from the first day in the 1 to 30 s reward-scarce environment (day 16, a, c) and the last day in this experiment (day 55, b, d). Note that on day 16, in the offer zone, mice made enter vs skip decisions without factoring in cued offer cost but still displayed biases to enter depending on the ordinal ranking of each restaurant. In contrast, by day 55, experienced mice made enter vs skip decisions in the offer zone not only based on the ordinal ranking of each restaurant but also as a function of cued offer cost. The fact that experienced mice treat the same tones differently in each restaurant indicates mice integrate auditory information with spatial information. Horizontal dashed gray line indicates chance p(choice) at 0.5. Shaded error bars ± 1 SEM.

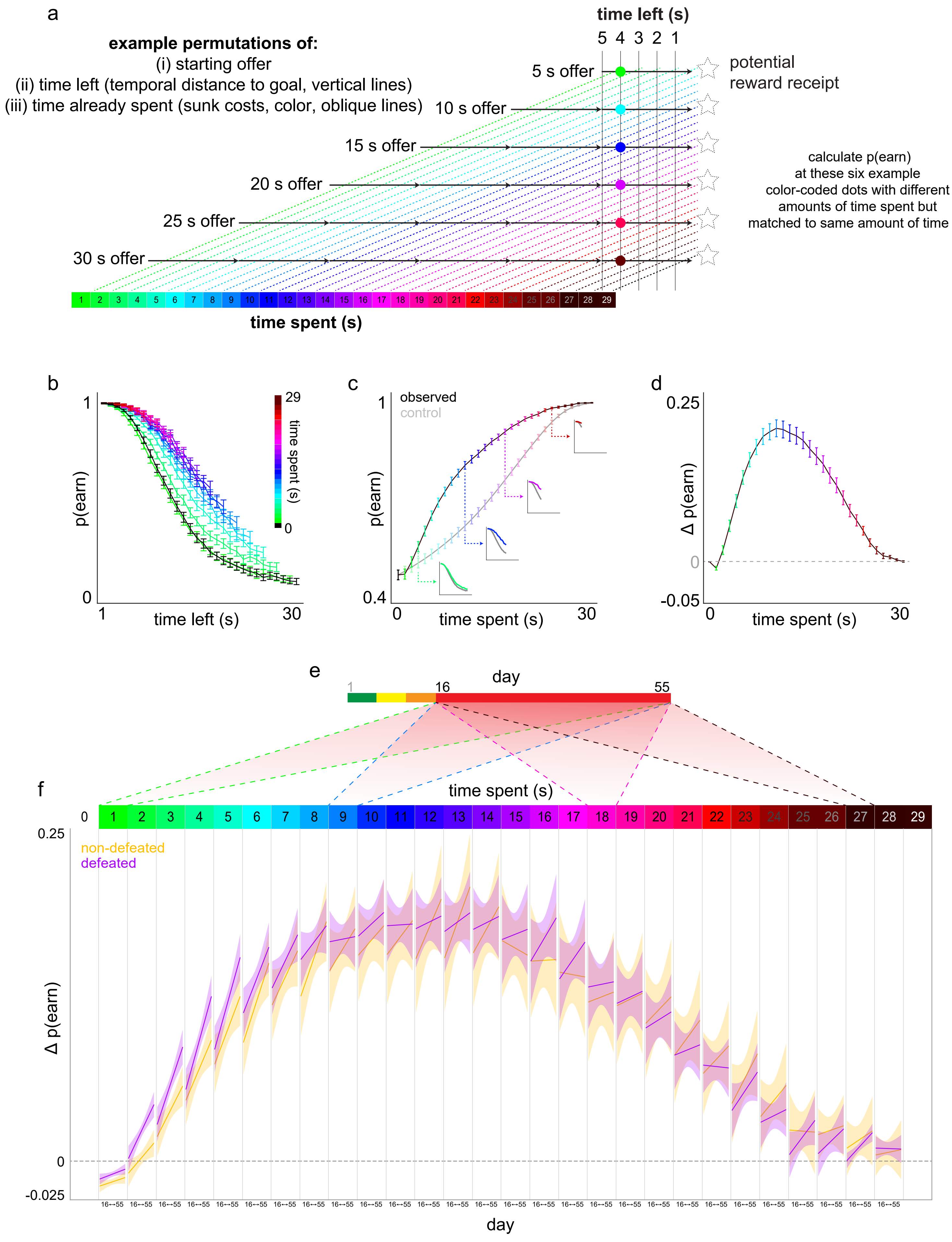
a
example permutations of:
(i) starting offer
(ii) time left (temporal distance to goal, vertical lines)
(iii) time already spent (sunk costs, color, oblique lines)
time left (s)
5 4 3 2 1
5 s offer
10 s offer
15 s offer
20 s offer
25 s offer
30 s offer
potential
reward receipt
calculate p(earn)
at these six example
color-coded dots with different
amounts of time spent but
matched to same amount of time left
time spent (s)
b
p(earn)
time left (s)
time spent (s)
c
observed
control
p(earn)
time spent (s)
d
Δ p(earn)
time spent (s)
e
day
f
time spent (s)
non-defeated
defeated
Δ p(earn)
day

**Supplementary Figure 5. Visual explanation of the sunk cost analysis with sensitivity rates across testing.** (a) All possible trial combinations of [time spent, time left] pairs can be compiled based on the randomly presented starting off costs and by using analysis points at progressively longer wait times during the countdown in the wait zone. Displayed here are 6 example starting offers (5, 10, 15, 20, 25, and 30 s offers, although this analysis extends to all 1 to 30 s offers). Colored diagonal dashed lines represent increasing amounts of time already spent waiting in the wait zone as animals progress toward earning a reward (dashed star shapes). 5 vertical solid blank lines reflect an example epoch of the last 5 s remaining in the countdown before potentially obtaining a reward common to all starting offer lengths. Using the 4 s remaining vertical solid black line as an example, the 6 color-coded dots indicate separate sunk cost conditions with increasing amounts of time already spent (colors) matched for the same temporal distance to the goal (4 s in this example). Of the trials in which the mouse had not yet quit by this point (i.e., trials "still live"), the probably of staying in the wait zone and earning a reward can be calculated at this analysis point. Sensitivity to sunk costs is represented by an increase in p(earn) as a function of time already waited, or an escalation of commitment. This effect is observed independent of and orthogonal to temporal distance to the goal because this analysis can be repeated at every instance of "time left" and a conserved effect of sunk costs on increasing p(earn) can be observed. By collapsing across the time left dimension and only focusing on the time spent (color) dimension, we can appreciate a change in p(earn) relative to the same amount of time left for all 0 s sunk cost conditions. (b-d) Data from the 1 to 30 s epoch across all 32 mice used to demonstrate analysis steps. (b) p(earn) in the wait zone calculated as a function of time left in the countdown (x-axis) and time spent (color). Note p(earn) increases as a function of time spent regardless of temporal distance to the goal. (c) p(earn) data from (b) collapsed across time left in order to capture envelope of sensitivity to sunk costs. Control analysis iteratively collapses the 0 s sunk condition (black curve) repeatedly matched to the same number of time left datapoints for a given sunk cost condition. Small inset graphs depict four examples from (b) collapsed into single datapoints in (c) with matching number of truncated datapoints in the 0 s sunk condition collapsed into the gray control curve. (d) The magnitude of sensitivity to sunk costs, Δp(earn), is calculated by subtracting the observed – control curves in (c). (e) This analysis was performed across the entire testing epoch (see main figure 4g-h) however here we focus just on the 1 to 30 s reward scarce environment (days 16 to 55) in order to display the change in p(earn) for each sunk cost condition across days of testing in (f). The x-axis in (f) reflects days 16 to 55 repeated for each sunk cost condition (four example shadow boxes from (e) show the reward-scarce epoch [red] compressed into each sunk cost condition). Linear fits across the entire day 16 to 55 epoch were performed for each sunk cost condition. Note that the rate at which mice acquire sensitivity to sunk costs across days of testing is enhanced in defeated mice particularly during the initial few seconds during re-evaluation change-of-mind decisions. Error bars ± 1 SEM. Shaded error bars 95% CI of linear fits.

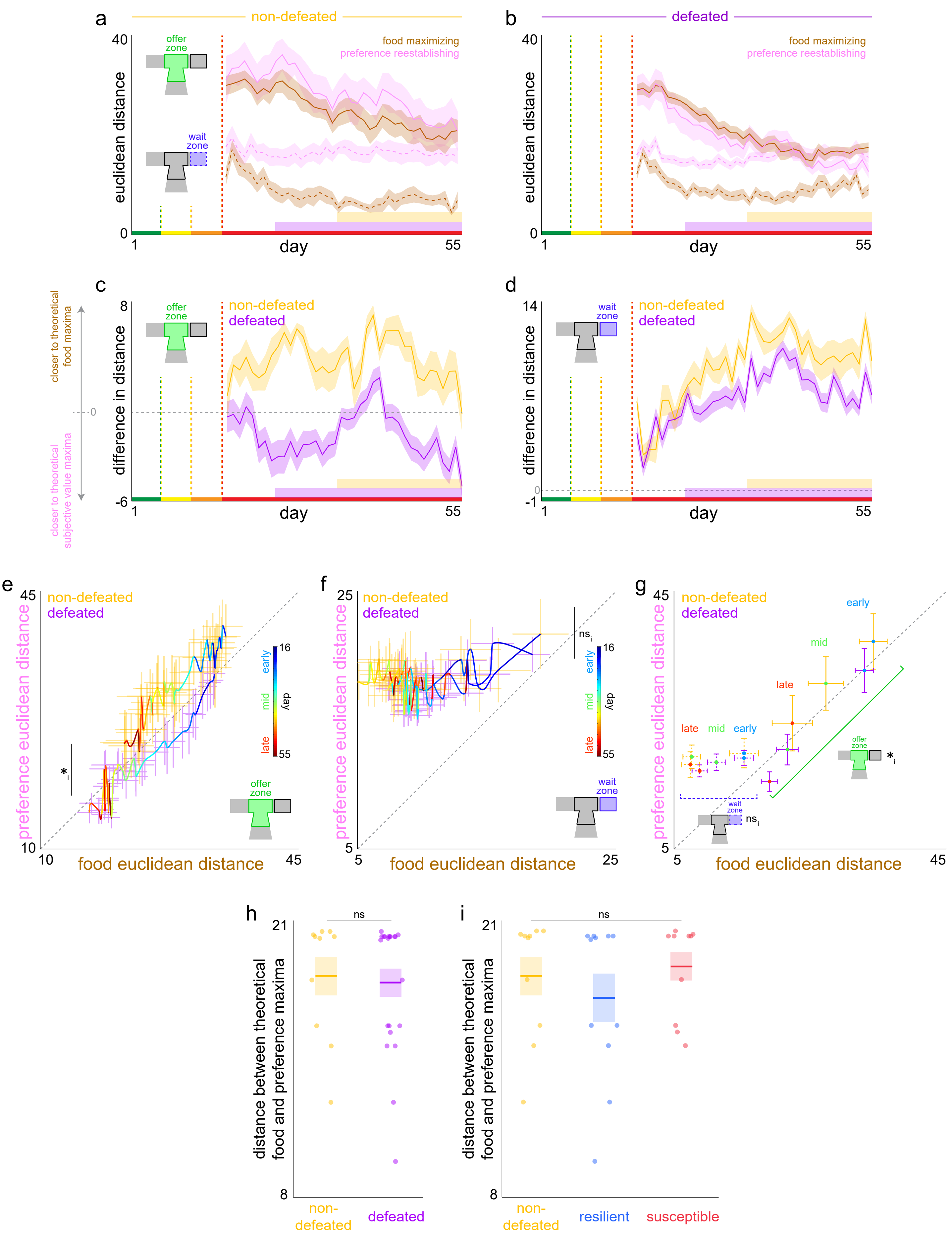

a
non-defeated
euclidean distance
offer zone
wait zone
food maximizing
preference reestablishing
day
b
defeated
c
difference in distance
closer to theoretical food maxima
closer to theoretical subjective value maxima
d
e
preference euclidean distance
food euclidean distance
early
mid
late
day
f
g
h
distance between theoretical food and preference maxima
ns
non-defeated
defeated
i
resilient
susceptible
